# Protocol Update: The Normative Modelling Paradigm for Computational Psychiatry

**DOI:** 10.64898/2026.02.17.706268

**Authors:** Augustijn A.A. de Boer, Johanna M.M. Bayer, Charlotte Fraza, Alice V. Chavanne, Barbora Rehak Buckova, Konstantinos Tsilimparis, Emin Serin, Antoine Bernas, Ramona Cirstian, Mariam Zabihi, Saige Rutherford, Abdool Al-Khaledi, Thomas Wolfers, Christian F. Beckmann, Seyed Mostafa Kia, Andre F. Marquand

## Abstract

Normative Modelling (‘brain growth charting’) is now a well-established method for computational psychiatry and involves charting centiles of variation across a population in terms of mappings between biology and behavior, providing statistical inferences at the level of the individual. These models have helped the field to move away from case-control analysis toward individual-level analysis. Correspondingly, normative modelling has now been applied to chart brain development and ageing in many populations and has been used to quantify individual deviations across various neurological and psychiatric conditions. This has been supported by large-scale models that are openly accessible for diverse brain imaging modalities. As normative modelling continues to grow, several recent methodological developments, such as non-Gaussian models, longitudinal models, and federated learning, have been implemented in different software tools, including the Predictive Clinical Neuroscience toolkit (PCNtoolkit). In this protocol update, we provide: (i) a revised overview of this methodological landscape; (ii) an update to our 2022 standardised analytical protocol for normative modelling of neuroimaging data, including options for federated and longitudinal normative models; (iii) practical guidance suited to both novice and experienced practitioners supported by open-source code examples implemented in the refactored version of PCNtoolkit; and (iv) updated models for cortical thickness, surface area, volumetric data, functional connectivity and diffusion-weighted imaging for use by the community.

## 1 Introduction

### 1.1 Development of the protocol

A major goal in the field of computational psychiatry is to develop reliable biomarkers for psychiatric disorders[1]. Traditional approaches based on case-control study designs have largely failed to achieve this and also fail to provide meaningful insight into psychiatric disorders because the rigid and categorical assumptions of the methodology do not match the continuous and heterogeneous nature of the underlying biology [2].

Normative Modelling (NM) has emerged as a mature statistical method for addressing these limitations by parsing heterogeneity in clinical cohorts[3]. The NM paradigm moves away from traditional case-control approaches by learning mappings between biology and behaviour. For example, mapping trajectories of healthy development – anatomical, neurological, behavioral – from a large reference dataset, and quantifies how individuals deviate from these norms. This is typically achieved by modelling biological or other response variables (e.g., regional brain volume) as a function of certain covariates (e.g., age, sex, total brain volume), thereby rendering the deviation space mostly independent of these variables and providing the ability to make statistical inferences at the level of the individual person.

The approach has been widely adopted in recent years. Large-scale normative models have been deployed [4–6] and applied to parse heterogeneity in several clinical applications [7, 8]. Currently, various methodological innovations have been proposed to improve the modelling of non-Gaussian data [9], longitudinal data [10–12], model comparison [13, 14], and federated learning [15]. These innovations allow researchers to more accurately estimate the mean and centiles of the population distribution, enable the tracking of deviation scores over time, compare models to find which one gives the most accurate deviation scores and update model parameters without sharing data.

The Predictive Clinical Neuroscience toolkit (PCNtoolkit) is an open-source Python package for normative modelling, which was developed to fulfill two critical needs. First, to provide a flexible and user-friendly software package for normative modelling, specifically tailored to neuroimaging data, and second, to ease reproducibility efforts. The initial distributions of the PCNtoolkit were published under the name `nispat`, with the first version of PCNtoolkit being released as version `0.15` in 2020. A protocol was published for version `0.20` of the PCNtoolkit in 2022 [16], and continued development led to the last release within this generation, version `0.35` in April 2025. Ongoing method development and an increased adoption of the NM paradigm demanded a major refactoring of the PCNtoolkit to keep it usable and maintainable. This led to the PCNtoolkit version `1.x.x`, with an easier interface to familiar NM techniques, refinement of model estimation and inference techniques in addition to new features such as longitudinal modelling and federated pipelines.

### 1.2 Overview of the procedure

The present protocol provides an update to the 2022 protocol [16] and: (i) provides an overview of the recommended workflow for normative modelling, illustrated with examples of the main methods implemented in the latest PCNtoolkit version `1.x.x`, and (ii) accommodates the updated syntax and also showcases new features, including new estimation algorithms for modelling non-Gaussian distributions [9] and longitudinal data [12]. In addition, we also have updated our existing large scale models for cortical thickness, surface area, subcortical volume, functional connectivity and fractional anisotropy in PCNtoolkit `1.x.x` and make these models available for use for the community. These models supersede our prior published reference models [4, 17, 18] and in many cases also offer expanded sample size and improved age range coverage.

In more detail, this protocol provides updated code examples for the methods covered in the 2022 protocol, and extends the protocol by covering longitudinal normative modelling, hierarchical Bayesian regression (HBR), model comparison using expected log pointwise predictive density (ELPD) [19], data harmonization, post-hoc-analysis suggestions, and federated learning methods; transferring, extending, and merging models. See figure 1 for an overview of the protocol.

**Fig. 1:**
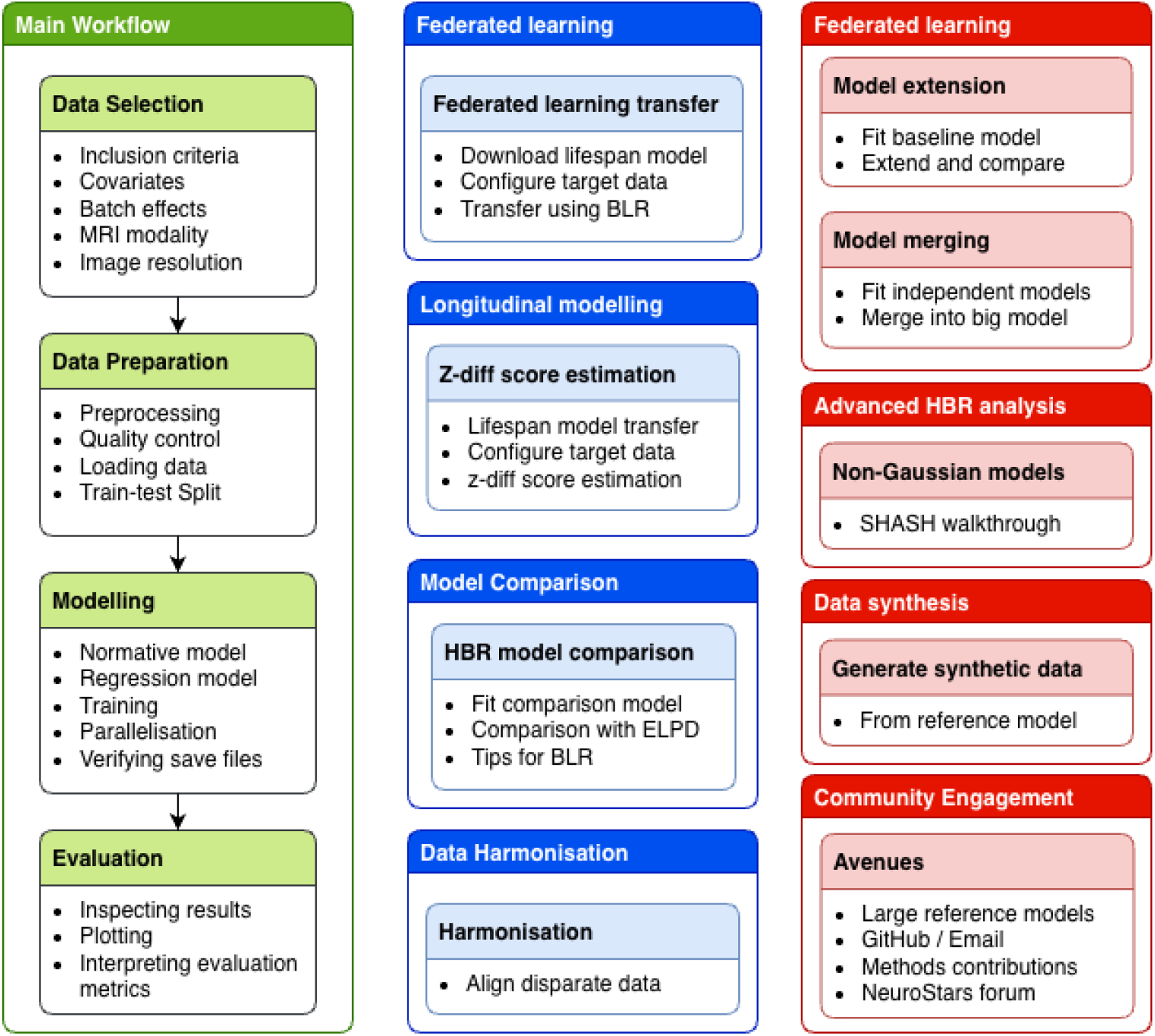
Overview of the protocol. The main workflow is shown in green. Optional secondary workflows that are also covered in the protocol paper are shown in blue. Additional workflows and other resources are shown in red (for example corresponding to additional notebooks available in the associated code repository).

**Fig. 2:**
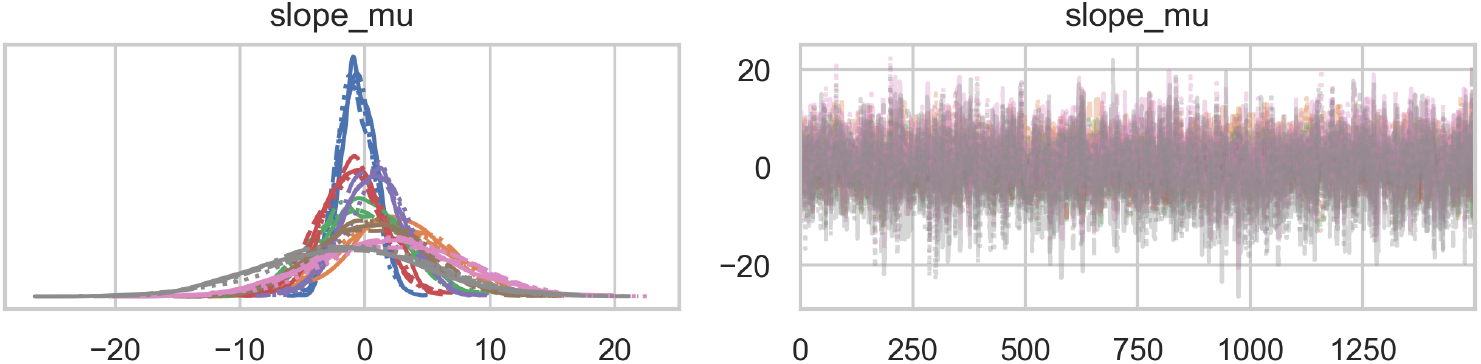
Traceplots for the covariate slope parameters (`slope mu`) from the Normal likelihood model. Each color represents one of the MCMC chains. The dense, well-mixed traces indicate stable posterior exploration and full chain convergence.

Rather than a comprehensive PCNtoolkit guide, this protocol provides general recommendations to brain charting through normative modelling, accompanied by code examples written for the PCNtoolkit version `1.x.x`. For a complete reference to the PCNtoolkit, see the [PCNtoolkit documentation](https://pcntoolkit.readthedocs.io/en/latest/). Despite the focus of this protocol on PCNtoolkit, the general principles and workflows are also applicable to other software implementations [5, 6, 20, 21].

The protocol is structured as follows: the core workflow outlined in the Procedure section covers the basic sequential process that can be followed to fit a normative model from scratch with PCNtoolkit. In addition, we provide a set of optional secondary workflows that can be applied in any order, but require a fitted model (i.e., completed main workflow). These optional workflows require one or more pre-estimated models, and in some cases an independent training dataset.

In the main workflow, the reader is guided through a complete workflow of creating a set of normative models with PCNtoolkit. First, for completeness, we provide a summary of the NM paradigm, what the goals are, and how the NM paradigm can be applied to neuroimaging data. Then, we walk through the installation and verification of the PCNtoolkit. This is followed by a step-by-step guide through the process of creating a normative model. The user is guided through the selection and filtering of a dataset on the basis of clinical and demographic information, data missingness, research question, and data modality. Then, we discuss Quality Control (QC) procedures and demonstrate how to use the PCNtoolkit to perform some basic data preprocessing steps, such as outlier detection and cleaning, missingness detection and handling, train-test splitting, and data scaling. We select and configure the algorithm and model parameters, which depend on the data regime, model requirements, and the research question. We fit the model on the selected and prepared dataset, and discuss the fitting of NMs in parallel on a compute cluster, and the evaluation of models. We then verify the output and discuss the interpretation of the resulting plots, results, and model evaluation metrics.

In the optional federated learning workflow we demonstrate how to transfer a large scale reference model to a clinical sample. We also provide more extensive examples of federated learning in the associated code repository, which provides more detailed examples of the three strategies for federated learning that are implemented in the PCNtoolkit: i) model transfer, which allows for transferring a reference model to a new local dataset (resulting in a smaller local model which can be used on the local data), ii) model extension, which allows for extending a reference model with data from a new dataset (resulting in a larger reference model which can be used on both the new and the original dataset; and iii) model merging, which allows for merging two or more fitted models into a single model. We discuss the differences between these strategies and when to use each one. In the longitudinal modelling workflow, we build on this model transfer technique to illustrate longitudinal NM using Z-diff scores, which statistically quantify change over time on the basis of a large-scale cross-sectional model. In the model comparison workflow, we illustrate HBR model comparison using the expected log pointwise predictive density (ELPD), and how to use this framework for model comparison. In the final optional workflow, we show an example of how to use normative models for data harmonization.

We provide background material for these workflows in the in the next sections, although we refer the reader to the primary publications for full details (see e.g. key publications below). Finally, we also provide a number of additional workflows in the associated repository code repository that cover additional functionality beyond the scope of this article. This includes additional federated learning examples mentioned above, a didactic overview of non-Gaussian modelling and an example of data synthesis.

### 1.3 Applications

Normative modelling is a general framework for deriving subject-level deviation scores from a reference population, and as a consequence, it can be applied to a wide range of (medical) data and research questions. Here we list a few:

1. Derive a normative model from a reference population of healthy individuals, and draw further conclusions about the population based on the estimated model parameters and deviation scores. For example, in Gaiser et al. [22], a large-scale normative model is fit to the cerebellum to understand cerebellar development across the adolescent period.
2. Transfer a reference NM trained on a large, healthy population to a new, smaller local dataset from a different population, and use it to derive biomarkers from the local clinical samples, for example, as is done in the context of psychosis in Worker et al. [23].
3. Use normative models fit to cross-sectional data to identify centile crossings in longitudinal data, as is done in the context of schizophrenia in Rehak Buckova et al. [11] and in dementia in Bayer et al. [12].
4. Use z-scores as features in downstream analysis, for example, as features in supervised [17] or unsupervised [24] machine learning models.

### 1.4 Comparison to other methods

When NM is viewed as a framework for deriving centiles of variation from a reference population, a wide range of statistical methods can serve as its foundation. For example, the original Gaussian processes (GP) approach for NM [3, 25], but also other algorithms can be used, including Bayesian Linear Regression (BLR) [13], Hierarchical Bayesian Regression (HBR) [26, 27], generalised additive models of location, scale and shape (GAMLSS) [5, 28] and fractional polynomial regression [6]. Future developments may expand the range of methods that can be seen as forms of NM even further. We will not compare these specific methods here because all methods are highly flexible, depending on different parameterisations (e.g., the specific distributional form for the centiles, underlying basis for modelling non-linearity, etc.) and whether the model is probabilistic or not. These modelling choices are more important than the particular method chosen, and also make direct comparison difficult. Rather, we believe that the most important aspect of NM is not the underlying regression method itself, but the framework that it provides for analyzing deviation scores from a reference population. The important feature of normative modelling is that it estimates centiles of variation in addition to the mean. In contrast, a regression model that is fit to data but does not provide centiles of variation, such as a simple linear regression model, can be seen as an approximation to NM when viewed in this light. However, such simple methods do not properly account for different variance components [7] and may not provide a good fit of the estimated centiles to the underlying data, which can bias downstream inferences [9, 29]. Moreover, it is increasingly recognised that the choice of reference cohort is of equal importance to the choice of algorithm, because of the potential for introducing demographic or racial bias [30, 31].

There are several other toolboxes available for neuroimaging data. PCNportal [20] is based on PCNtoolkit and provides a code-free repository for pre-estimated structural, functional and diffusion based models that is accessible via a browser. Bethlehem et al [5] introduced an interactive R shiny application that also provides browser based access to pre-estimated structural models fit using GAMLSS. Ge et al [6] introduced CentileBrain which includes structural models fit using Gaussian fractional polynomial regression. Finally, Little et al [21] introduced a software toolbox and models for normative modelling also based on GAMLSS. We focus here on PCNtoolkit as it provides a comprehensive python-based library of tools for fitting and evaluating models built either from scratch or in a federated learning context. However, many of the principles outlined in this can also be implemented using custom code in other platforms, for example using GAMLSS packages implemented in R [28].

### 1.5 Experimental Design

Several important design decisions should be considered before applying this protocol. The most fundamental choices in designing a normative modelling analytic strategy are the choice of response variables, covariates and batch effects, the choice of reference cohort and data partitioning strategy and the choice of modelling approach (including the choice of algorithm and likelihood function).

#### 1.5.1 Basic Configuration

##### Covariates, batch effects and response variable selection

Normative models fit using PCNtoolkit version `1.x.x` consist of one or more regressions of response variables on the selected covariates, together with a mechanism to handle batch effects, the details of which depend on the model type. The choice of covariates and batch-effect variables depends on the research question, but they should generally include all factors known to influence the measurements and be available in all datasets, such as age, sex or site. Batch effects are used to group subjects together in the same way that random effects are used in classical multilevel modelling. Typically, batch effects are used to model nuisance variation such as site or scanner effects, but they could also be applied to model other features such as sex, ancestry or imaging field strength. Whilst batch effects can also be encoded as dummy encoded covariates, this is less flexible and is equivalent to a fixed intercept model (i.e., only allowing shifts in the intercept). In contrast, using batch effects allows the model to estimate separate slope, intercept and basis function parameters for each site with partial pooling across levels of the batch effects (see [15, 26]).

The covariates are typically continuous variables, like age or height, but can also be numerically encoded categorical variables, for example, using one-hot-encoding. The batch effect dimensions are categorical variables, for example, scan site or scanner brand/model. The response variables are continuous variables, like brain volumes or brain activity patterns. All these variables must be available for all subjects in the reference set. Many normative models predict brain-derived measures (response variables) based on age and sex (covariates), whilst accounting for site (e.g., using batch effects), this is not the only way such models can be employed. For example, in [3] a normative model was constructed to predict reward-related brain activity on the basis of delay discounting scores, which measure individual variability in reward sensitivity and are sometimes taken as a proxy for impulsivity. In [32], a normative model was constructed to predict brain structure on the basis of measures of adversity.

When selecting covariates, care must be taken when including variables that are highly correlated with one another, as multicollinearity can inflate parameter uncertainty and make it difficult to interpret the contribution of individual covariates. However, this is principally relevant for applications where parameter interpretation is important.

##### Reference cohort selection

The reference cohort and data partitioning strategy should also be carefully considered (i.e., how to divide the available data into training and test sets), which directly influences how the deviations from such a model should be interpreted. Whilst these aspects have been covered elsewhere [7, 8, 16], in brief, a large reference – or normative – set is required to train a well-calibrated normative model. What can be considered a reference set depends on the research question. If the goal is to understand variation in a given population, the reference set could be sampled from this population. If the goal is to parse heterogeneity within a clinical cohort, one could choose either a healthy or population reference or choose to use a set of participants with that particular condition as the reference set. Regardless, the reference cohort should be chosen to be as representative as possible of the population of study interest. If a healthy cohort is chosen, then deviations should be understood with respect to this cohort. Alternatively if a population-based cohort could be used (i.e., including individuals with brain disorders), the prevalence of each subgroup in the reference cohort should be taken into consideration when interpreting the target cohort, where the prevalence may be different. In many cases (for example, for rare conditions), it may be desirable to ensure that a sufficient number of individuals having the phenotype of interest are retained in the test set to maximise the possibility of detecting downstream effects. In other words, the training and test sets should be stratified for the clinical labels.

A generalizable normative model should also incorporate two types of uncertainty: epistemic and aleatoric [7]. Epistemic uncertainty refers to the uncertainty in our knowledge (e.g. modelling uncertainty or uncertainty in the parameter estimates). Epistemic uncertainty decreases with more data. In contrast, aleatoric uncertainty arises from the natural variation in the population (e.g. inter-individual differences) and does not decrease with more data. The latter is what we aim to capture with our normative models; to model it as accurately as possible, the former should be minimized. Reducing epistemic uncertainty is a major reason why it is desirable to have a large reference data set.

It is also important to consider the range of the covariates in the choice of reference cohort [33]. For example, since many datasets have a limited age range, which can introduce collinearity with batch or site effects, it is desirable to have overlap (i.e., multiple datasets spanning the same age range) to assist in disentangling these effects.

##### Data preparation, quality control, and train-test split

Preparing the data for modelling involves a sequence of preprocessing steps designed to ensure data quality and reproducibility. Note that standard neuroimaging preprocessing steps such as spatial normalisation and/or cortical surface reconstruction should be applied first. All subjects in a normative modelling analysis need to be in the same space.

Steps specific to normative modelling include handling missing values and outliers, combining multiple data sources, splitting the data into training and test sets, and rescaling covariates and response variables (e.g., via standardisation). All procedures are implemented in the PCNtoolkit, with the `NormData` class providing dedicated functionality for data cleaning, harmonization, and preparation.

PCNtoolkit automatically excludes subjects with missing values and identifies outliers based on feature-wise Z-scores. The stringency of this filter is controlled via the `z threshold` argument. Note that this is done separately for each phenotype (to minimise loss of data when large numbers of phenotypes are considered). In this tutorial, we set *z*_threshold_ = 10 to illustrate the procedure while avoiding unnecessary data loss; however, for practical analyses, a different value may be selected to balance sensitivity and specificity. However, care needs to be taken not to be overly aggressive with outlier removal as this may result in truncating the empirical distribution, thereby making it difficult to model with a continuous distribution. While the choice of the Z threshold depends on the characteristics of the dataset used for normative modelling, there are some practical considerations that may be generally applied. Removing outliers relative to the model will always come at the cost of potentially removing meaningful variance. We recommend not going below a threshold of |*Z*| *>* 3, as removing too many subjects can eliminate valid biological variation and truncate the distribution. Applying a threshold of |*Z*| *>* 3 or 5 may be appropriate when modelling most datasets (especially when these are known to be affected by artefacts), a more lenient threshold (e.g. |*Z*| *<* 7) may be preferable to preserve heterogeneity. If a customized approach to missingness or outlier handling is desired, data may also be preprocessed externally before being passed to PCNtoolkit.

The train–test split is performed using the `train_test_split` function from `scikit-learn`, with automatic stratification on batch-effect variables (e.g., `site` and `sex`) to maintain balanced distributions and to avoid demographic or acquisition drift between subsets. Data scaling is applied internally, and the corresponding scaling coefficients are stored within the fitted model object to ensure that identical transformations are applied to all future data during prediction, or when transferring the model.

The reference set is divided into a training set used for model fitting and a test set reserved for model evaluation. Assessing the model on independent data is critical to ensure accurate estimates of generalisabilty. If the goal is to provide a reference model for new data, the model can be refit to the whole dataset once generalisability has been established. A trade-off exists between model accuracy and the reliability of performance estimates: a larger training set improves model precision, whereas a larger test set yields more stable validation metrics. We recommend an 80/20, 70/30 or 50/50 train–test ratio, depending on dataset size.

To obtain unbiased evaluation metrics, the test set should be statistically similar to the training set (i.e. be drawn from the same population). This can be achieved in PCNtoolkit by stratification of the train–test split, which ensures that the distributions of key batch-effect variables (e.g., site, sex) remain consistent across subsets. However, it may also be of interest to study differences between test sets having different characteristics. For example, a test set containing individuals with neurodegenerative or psychiatric disorders can often have poorer fit metrics than a test set that is comparable to the training set. This is expected considering that differences between the cohorts are then reflected in the fit metrics.

##### Algorithm selection

The choice of algorithm should also be carefully considered. PCNtoolkit, `1.x.x` currently supports two algorithms, BLR (with or without likelihood warping) and HBR. These have many similarities: they both are either linear or employ a b-spline basis expansion to model non-linearities. In addition, both models support Gaussian or non-Gaussian noise models via the sin-arcsinh (SHASH) transformation (see the next section or [9, 13] for details). However, they also have differences. The most important difference from a modelling perspective is that BLR specifies a linear model for the data uses an empirical Bayes approach to optimise model hyperparmeters governing the variance and shape of the distribution, whereas HBR assumes a hierarchical linear generative process for the data and performs full posterior estimation for all model parameters via Markov Chain Monte Carlo (MCMC). This means that BLR is considerably faster (and is therefore preferred when many models need to be estimated, for example for voxel-wise data) but does not fully account for the uncertainty in model parameters. In contrast, HBR provides greater modelling flexiblity for batch effects and also support for the Beta likelihood (which is not available for BLR), but is considerably slower to fit.

Since HBR is more complex, we focus on this model for this protocol, but an example configuration for BLR models is provided in the companion code repository: https://github.com/predictive-clinical-neuroscience/pu25_code/blob/main/notebooks/main_workflow.ipynb, and the entire BLR workflow used to fit the lifespan models used in the transfer learning section of the protocol is provided here: https://github.com/predictive-clinical-neuroscience/braincharts_pu25/. In addition, the optional federated learning and longitudinal workflows are also based on BLR.

#### 1.5.2 Implementation details in PCNtoolkit

We next describe some of the core components in PCNtoolkit that are necessary to understand the workflow provided in the protocol outlined below.

##### The NormativeModel class

The central class in PCNtoolkit is the `NormativeModel` class. It has all the functions that are required to build and use a normative model. To construct a `NormativeModel` object, the user needs to provide a template regression model, which is copied and fit for each feature in the reference set.

##### The RegressionModel class

The current implementation of PCNtoolkit supports the creation of univariate normative models. This means that each `NormativeModel` object contains a collection of `RegressionModel` objects, each of which provides the regression functionality that underpins the normative model estimation. Concretely, for each response variable, the `NormativeModel` class makes a copy of a template regression and fits it to the response variable. This template is what the user should provide when creating a `NormativeModel`. There are currently two `RegressionModel` variants instantiated in PCNtoolkit: BLR and HBR, and both come with their strengths and weaknesses as oultlined above.

##### Parallel fitting with the Runner class

For large datasets (either in terms of the number of samples and response variables), fitting models sequentially can be quite time-consuming. In case of access to a high-performance computing cluster running standard workload managers such as Torque or Slurm, operations such as `fit_predict()` can be parallelized by using the `Runner` class. The Runner class will allow running the same tasks as in the previous step, but in a distributed fashion. This is a little more involved than in the previous step. For example, you need to specify the Python executable and time and memory limits for each job. You can also specify the number of batches into which to split the workload.

##### Standardization

In order to ensure that there is no data leakage between training and test sets, PCNtoolkit automatically stores the scaling coefficients used during feature scaling parameters within the fit model object. Note that these are derived from the training set only and can be applied to the test set. This ensures that all subsequent predictions and transferred models apply the same transformations to incoming data, eliminating manual preprocessing and guaranteeing consistent scaling across datasets during model development and deployment stages.

##### Configuring likelihood function

A crucial decision in normative modelling is to determine the appropriate likelihood function to model the centiles of variaion in the data. PCNtoolkit currently supports three likelihood functions: Normal, sin-arcsinh (SHASH), and Beta likelihoods, each suited to different data characteristics of the data. Each likelihood has interpretable parameters, which can each be modeled hierarchically to capture structured variation across covariates and batch effects. This topic is covered in detail elsewhere [9, 34], however for completeness a brief overview is given here.

The Normal likelihood is used for continuous, symmetric and approximately Gaussian response variables (*y*), and is parameterised as:

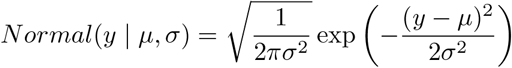

where *µ* and *σ* describe the mean and standard deviation.

The SHASH likelihood is suited for continuous data that are positively or negatively skewed and/or platykurtotic or leptokurtic.

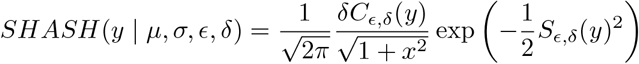

Where *S*_*δ,ϵ*_(*y*) = sinh *ϵ* + *δ* sinh^−1^(*y*) and *C*_*δ,ϵ*_(*y*) = cosh *ϵ* + *δ* sinh^−1^(*y*). This distribution is governed by four parameters, namely *µ* and *σ* (which are handled implicitly by rescaling the random variable *y*) and correspond to the mean and standard deviation. In addition, the variables *δ* and *ϵ* model the kurtosis and skew respectively. In PCNtoolkit, the SHASH distribution is supported in both BLR (via likelihood warping) and HBR.

The Beta likelihood is suited for data that are confined to a given interval (e.g. bounded between zero and one). The probability density function for the Beta likelihood is:

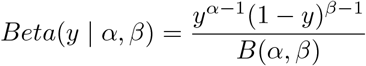

Where *B* is the beta function. The Beta likelihood is parameterised by *α* and *β* where the mean is *α/*(*α* + *β*) and the variance is *αβ/*((*α* + *β*)^2^(*α* + *β* + 1)). In PCNtoolkit, the Beta likelihood is only available for HBR.

For HBR, once a likelihood is chosen, it is necessary to define priors over its parameters, which may include both fixed and random effects. These are specified using the `make_prior` function. Notably, parameters can vary linearly with covariates, include random (batch) effects, or remain fixed across all observations. Random-effect priors are implemented in a non-centered hierarchical form (reparameterized) to stabilize sampling and avoid funnel pathologies during inference. See [9, 34] for more details.

An additional step for HBR is to assess model convergence, that is to evaluate whether the Markov chains used for inference have converged to a stationary distribution. There are several standard statistical metrics to assess this [35] which are based on the intuition that different chains should yield solutions with the same distributional properties.

##### Model Assessment

A summary of the main model fit metrics in PCNtoolkit is provided in Table 1. In general, a lower root/standardised mean squared error (RMSE/SMSE), mean standardised log loss (MSLL) and mean absolute centile error (MACE) values, and higher explained variance (EXPV) indicates better overall fit. However, it is important to recognise that the interpretation and evaluation of the fit indices are dataset-dependent. Some metrics (e.g. RMSE) depend on the scale of the data (e.g. implicitly expressed in units of cortical thickness). The SMSE overcomes this problem by removing the scale of the data such that a value of one is the performance of a dummy baseline model that always predicts the mean of the training set. Therefore anything less than this can be considered better. Even for metrics that do not directly depend on the scale, such as EXPV, it is difficult to provide general guidelines as to what can be considered good performance, because this is still dependent on the particular dataset under comparison. For instance, the EXPV is usually lower for a dataset having a narrow age range (which makes it difficult for age to explain much variance), relative to a dataset which spans the entire lifespan (where there is more variance to explain). Nevertheless, we provide some useful rules of thumb for assessing model fit: the EXPV should be positive for a model that fits well, but can be close to zero or negative for poorly fitting models. Unlike the classical (in-sample) statistical notion, the EXPV is estimated out of sample and therefore can be negative which reflects the fact that regression models can become arbitrarily bad for independent data.

**Table 1:** Summary of evaluation metrics 43.

|  | Full Name | What it measures | Interpretation | Mean-based |
| --- | --- | --- | --- | --- |
| EXPV | Explained Variance | Proportion of variance in the data explained by the model (similar to $R^2$, but can differ). | Closer to **1** = better (0 means no variance explained, negative means worse than mean predictor) | Yes |
| MACE | Mean Absolute Centile Error | Average absolute deviation of predictions from targets expressed in percentiles. | Closer to **0** = better (0 means perfect percentile alignment, higher means worse calibration). | No |
| MAPE | Mean Absolute Percentage Error | Average of absolute percentage errors between prediction and true values. | Closer to **0%** = better (can blow up if actual values are close to 0). | No |
| MSLL | Mean Standardized Log Loss | Compares model log-likelihood to a Gaussian baseline. | **Lower** is better (0 = same as baseline, negative = better than baseline, positive = worse). | Yes |
| MLL | Mean Log-Loss (previously named Negative Log-Likelihood (NLL)) | Measures how unlikely the observed data are under the model predictions. | **Lower** is better (0 = perfect likelihood, large positive = poor fit). | No |
| R2 | Coefficient of Determination ($R^2$) | Fraction of variance in target explained by the model. | Closer to **1** = better (0 means no variance explained, negative means worse than mean predictor). | Yes |
| RMSE | Root Mean Squared Error | Square root of the mean squared difference between predictions and true values. | Closer to **0** = better (represents error in original units of the target). | Yes |
| Rho | Pearson Correlation Coefficient ($\rho$) | Linear correlation between predicted and true values. | Closer to **1** or **-1** = stronger correlation (0 = no linear correlation). | No |
| Rho.p | p-value for Pearson's $\rho$ | Statistical significance of the correlation. | Closer to **0** = stronger evidence of correlation (usually $p < 0.05$ is significant). | No |
| SMSE | Standardized Mean Squared Error | MSE normalized by the variance of the true data (relative measure of fit). | Closer to **0** = better (1 = as bad as predicting the mean, $>1$ = worse than mean predictor). | Yes |
| ShapiroW | Shapiro-Wilk Test Statistic | Tests normality of residuals (how Gaussian the errors are). | Values close to **1** = residuals look normal. Low values with small p-value indicate non-normality. | No |
| Skewness | Skewness | Measures asymmetry (i.e., whether one tail is longer than the other) of the Z-score distribution relative to a normal distribution. | Closer to **0** = better ($> 0$ indicates the model tends to underpredict, $< 0$ indicates the model tends to overpredict). | No |
| Kurtosis | Excess Kurtosis | Measures how heavy the tails of the Z-score distribution are relative to a normal distribution. | Closer to **0** = better ($> 0$ indicates heavier tails/more outliers, $< 0$ indicates lighter tails/fewer outliers). | No |

**Table 2:** Aggregated posterior diagnostics across all parameters (74 total). Values are shown as median [IQR].

| Metric | Median | IQR |
| --- | --- | --- |
| $\hat{R}$ | 1.00 | 1.00–1.01 |
| ESS<sub>bulk</sub> | 4513 | 2176–5502 |
| ESS<sub>tail</sub> | 3789 | 2533–4454 |
| Posterior mean | -0.11 | [-0.61, 0.35] |
| Posterior SD | 0.42 | [0.24, 0.92] |
| Monte Carlo SE (mean) | 0.006 | [0.003, 0.026] |

It is crucial to recognise that the RMSE, SMSE and EXPV are only dependent on the mean prediction and are insensitive to calibration (i.e. the match between the data and the shape of the predictive distribution). The mean standardised log loss (MSLL) improves on this slightly by also accounting for the predictive variance of the data, but it still does not account for the shape. We expect MSLL to be negative, with positive values indicating a poor model fit, and values closer to zero implying only weak improvement over the dummy baseline model described above.

In fact, in normative modeling calibration is nearly always of far more importance than predicting the mean in that a model having good calibration, but poor (even zero) EXPV can still be highly useful, but the converse is not true. This reflects the purpose of normative modeling: to accurately estimate the conditional distribution of each feature rather than only its central tendency. The best way to assess calibration is to examine the centile and QQ plots (e.g. Fig. 3). Alternatively, one can examine statistics such as the skew and kurtosis of the resulting Z-scores (see [13]) or other statistics such as the MACE and Shapiro-W statistics, which are sensitive to the shape of the distribution.

**Fig. 3:**
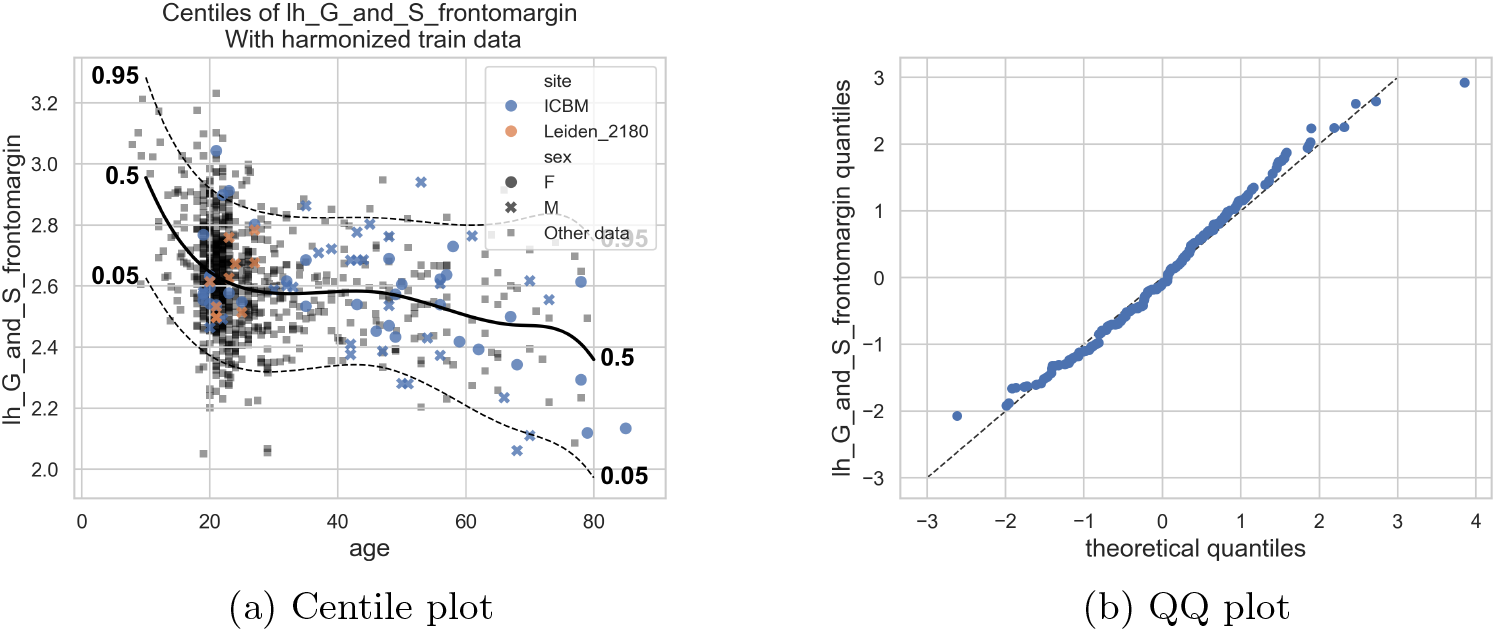
Visualization of normative model calibration for a representative region of interest. **(a)** Predicted centiles (5th, 50th, 95th) across age, with observed data overlaid. Smooth median and outer centile alignment indicate a good functional fit across covariates. **(b)** Quantile–quantile (QQ) plot comparing empirical versus theoretical quantiles of standardized residuals. Alignment with the identity line indicates accurate predictive variance; curvature suggests over- or under-dispersion. For a detailed interpretation of QQ plots in normative modelling, see [28].

Qualitative assessment of calibration is performed using PCNtoolkit’s built-in plotting utilities. `plot_centiles` visualizes predicted centile trajectories across covariates, while `plot_qq` compares empirical and theoretical quantiles of standardized residuals (Z-scores). Together, these provide complementary views of functional and distributional model fit.

#### 1.5.3 Federated Learning

A key feature of PCNtoolkit is federated learning, as described in detail in [15, 26]. Concrete examples of this workflow are provided in the optional federated learning workflow below and as notebooks in the associated code repository. In brief, PCNtoolkit provides three strategies for federated normative modelling, i.e., estimating normative models on decentralized data: `transfer`, `extend`, and `merge`. Each strategy provides a distinct mechanism for reusing information from an existing reference model without requiring access to the original data. This is critical for privacy-sensitive data such as clinical neuroimaging data.

Model transfer is a strategy for adapting a pre-trained reference model to a new site by re-estimating parameters using local data while constraining the solution with prior information derived from the reference model. In fact, it distills useful information from bigger reference model into a smaller local model. In the context of HBR, this is implemented by initializing the local model with informative priors corresponding to the posterior distributions of the reference model parameters. For BLR, the hyperparameters are adjusted using a post-hoc fitting procedure. In either case, the effect is to transfer information from the reference model to the smaller target dataset, effectively regularizing the estimates and potentially improving both centile estimation and generalizability. The target dataset is then used to update these parameters via Bayesian inference. Whilst this approach is usually the most efficient way deploy a large scale model to a smaller sample, model transfer may be problematic if the smaller target dataset has a very different distribution (e.g., non-overlapping covariate ranges or strong site effects) from the data in the training set. Moreover, model transfer does not preserve the reference model parameters for the original model but only the adapted parameters to the local dataset.

Model extension is more involved. This allows a given reference model to be augmented (‘extended’) with additional samples without the need to access the reference dataset. This is achieved by exploiting the generative nature of the model: generated (synthetic) samples are first drawn from the posterior predictive distribution of the existing model, representing the learned population-level variation. These synthetic data are then combined with the real data from the new site, and a full model is reestimated on the combined dataset. Compared with transfer, model extension retains information from earlier datasets (via the model parameters). Hence, it is a useful strategy in sequential federated learning scenario in which the model is passed through local sites one by one and it is repeatedly extended to derive a global reference model.

Model merging combines independently trained normative models into a single unified model. Under the hood, each model is first used to generate synthetic data from its learned distribution (via posterior predictive sampling). The resulting synthetic datasets are then combined, and a new global model is fitted to this combined synthetic dataset. In comparison to the two previous methods, model merging is a way to integrate many existing models post hoc rather than adapting a single reference model.

#### 1.5.4 Longitudinal normative modelling

The goal of longitudinal modelling is to develop a statistical model to chart change over time. Whilst many normative models are estimated using longitudinal data, the models themselves are fundamentally cross-sectional in nature in that they involve estimating smooth curves to link the group-level centiles across different timepoints. In order to obtain true longitudinal inferences, additional steps are necessary. Most importantly, it is necessary to provide a statistical estimate of the magnitude of change between two or more consecutive z-score measurements derived from a normative model. In other words, longitudinal normative modelling seeks to determine whether a centile crossing has occurred for a given individual between two or more timepoints. Since individuals do not track centiles exactly as they move through time (e.g., due to measurement noise), it is essential to take longitudinal variance into account both when estimating the normative model and when evaluating change between time points, to obtain a reliable characterization of individual trajectories [12]. This requires longitudinal data.

The PCNtoolkit and supporting scripts allow for two ways of obtaining an estimate of the significance of longitudinal change: namely constructing ‘Z-diff’ scores that provide a statistical estimate of centile crossing between two timepoints based on BLR models [11] or using velocity models, which provide the ability to estimate ‘thrive lines’ [12] which can quantify centile crossings for larger numbers of timepoints. An alternative method for longitudinal data analysis is presented in Di Biase et al. [36]. Here we employ the framework proposed in [11], where we statistically quantify difference over time using a ‘Z-diff’ score. In more detail, this score quantifies the longitudinal change in a response variable relative to what would be expected based on healthy ageing, normalized by the uncertainty in that prediction:

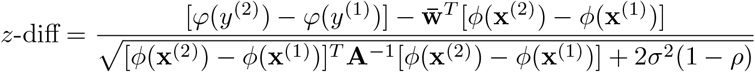

The denominator has two components, reflecting the uncertainties introduced in section 1.5, namely:

- **Epistemic uncertainty**: [*ϕ*(**x**^(2)^) − *ϕ*(**x**^(1)^)]^*T*^ **A**^−1^[*ϕ*(**x**^(2)^) − *ϕ*(**x**^(1)^)]. This reflects uncertainty in the model weights given the covariate change
- **Aleatoric uncertainty**: 2*σ*^2^(1 − *ρ*). This reflects the expected healthy between-visit variance, estimated empirically from longitudinal healthy controls

In this implementation, the epistemic uncertainty term is omitted from the denominator. This is because the posterior precision matrix **A** is estimated on the original training data and is not updated during transfer, making it unreliable for the new site. For the very large models used in this protocol (see ‘optional workflow’ below), the epistemic term is negligible compared to the aleatoric term (on the order of 10^−6^ vs 10^−2^), so this simplification has minimal impact on the results. The denominator therefore reduces to:

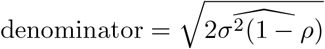

where 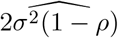 is estimated from the healthy controls as:

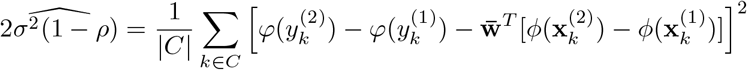

#### 1.5.5 Community Resources

##### Pre-estimated model dissemination

As outlined above and in order to maximise value to the community, we provide access to an updated set of lifespan reference models derived from tens of thousands of individuals aggregated from datasets that span the lifespan. These models supersede the cortical thickness and volumetric models presented in Rutherford et al. [4], fractional anisotropy measures derived from diffusion data [18] and other measures including surface area and functional connectivity presented in Rutherford et al. [17]. This is necessary because models estimated in early versions of PCNtoolkit are not compatible with PCNtoolkit `1.x.x`. ^1^These reference models are available for download via a notebook available in the associated code repository, at https://github.com/predictive-clinical-neuroscience/pu25_code/blob/main/notebooks/secondary_workflows/federated_learning_transfer.ipynb and as explained in the federated learning workflow below. We also welcome the contributions of new models from other groups via PCNPortal [20], our online collaborative normative modelling portal.

##### Methods contributions

We also welcome other contributions from the field, for instance new methods for normative modeling and different assessment metrics, as well as general improvements to the code. This was a major redesign consideration in PCNtoolkit `1.x.x` and is now much easier due to the object-oriented design. These can be submitted through GitHub issues or standard pull requests to the [PCNtoolkit GitHub repository](https://github.com/predictive-clinical-neuroscience/PCNtoolkit).

##### Additional information

Additional support can be found via the PCNtoolkit [documentation](https://pcntoolkit.readthedocs.io/en/stable/), via [GitHub](https://github.com/predictive-clinical-neuroscience/PCNtoolkit) issues or via the [Neurostars](https://neurostars.org/tag/pcntoolkit/1106) forum.

### 1.6 Regulatory Approvals

If the goal is to fit a normative model from scratch, the main regulatory consideration is permission to use the data used to fit the model, because in most instances it is necessary to aggregate cohorts to provide good coverage of the lifespan. This can either be performed using publicly available datasets or according to appropriate material transfer agreements. If the data cannot be shared, federated learning techniques can be employed to estimate normative models without the need to share data [15, 26]. Alternatively, if the goal is to transfer a pre-fit model to a new dataset, such approval is not necessary because this is done via the model coefficients, which are summary statistics that are no longer traceable to any individual participant. For details surrounding the public datasets we use to illustrate the protocol, please refer to the section 2.3 below.

### 1.7 Limitations

The principal limitation of this protocol is that in a document of this scope, it is not possible to demonstrate all the functionality of PCNtoolkit, nor provide a recipe that will work for all different kinds of modelling strategies, distributions or data types. This is an inherent to the expanding functionality of the normative modelling framework. However, we have selected a core workflow that provides a good basis for most analyses, plus optional secondary workflows that cover the most common aspects. On the associated code repository, we provide [additional workflows](https://github.com/predictive-clinical-neuroscience/pu25_code/tree/main/notebooks/additional_worflows) that cover more advanced functions. We also provide extensive [documentation](https://pcntoolkit.readthedocs.io/en/stable/) that can guide users toward additional functionality not covered here.

## 2 Materials

### 2.1 Hardware

Computing infrastructure required to run this protocol: a Mac, Linux or Windows with installed Windows Subsystem for Linux (WSL) computer with enough space to store the derived phenotypes from imaging data of the train and test set. In addition, an HPC infrastructure (Slurm or Torque) can be beneficial to parallelize fitting large numbers of normative models (e.g., for high-resolution data)

The code examples in this protocol have been tested on a MacBook Pro with an M3 Chip and 18GB of RAM. The dataset used in these experiments is rather small, so everything should be possible to run on any computer with a similar or lower configuration. Experiments using a larger dataset will require more RAM, and we recommend using a compute cluster for those cases (see Step 11(B) and Section 1.5.7. for more details about fitting models in parallel using the PCNtoolkit). The resources requested for the jobs in these experiments are always 16 GB of RAM and 4 cores.

Considering the available hard disk space, we recommend having at least 10 times the size of the dataset in free hard disk space. This is because the PCNtoolkit stores the predicted centiles (5 centiles per subject per feature), the Z-scores and the log-probability of the observed values (both 1 per subject per feature), which all together add up to 7 times the size of the original dataset. The saved model files and the plots, which are generated by the PCNtoolkit, also take up additional space. However, these functionalities can be disabled if the user desires to save disk space. A stable internet connection is also required to run the protocol.

### 2.2 Software

The protocol requires Python 3.12 and a few core scientific packages. Python can be installed from the official Python website (https://www.python.org/downloads/) or via Anaconda (https://www.anaconda.com/download). The main package used in this protocol is PCNtoolkit (current version 1.x.x), which can be installed via `pip` following the instructions at https://pcntoolkit.readthedocs.io/en/latest/pages/quickstart.html.

#### Software environment

In this manuscript, all analyses were performed in Python 3.12 and were tested with PCNtoolkit version 1.3.0. We recommend using this PCNtoolkit version or a later release. Older versions are not guaranteed to be compatible with this protocol. Additional core packages were: PyMC 5.25.1; ArviZ 0.22.0; PyTensor 2.31.7; NumPy 2.2.6; SciPy 1.16.1; scikit–learn 1.7.2; pandas 2.3.2; xarray 2025.9.0; xarray–einstants 0.9.1; matplotlib 3.10.6; h5py 3.14.0; h5netcdf 1.6.4; nutpie 0.15.2 (for accelerated sampling); and seaborn 0.13.2 (for figure styling).

Random seeds were fixed globally using `np.random.seed(42)` and `pymc.set_tt_rng(42)` to ensure bitwise-reproducible sampling, data splitting, and posterior predictive simulations. All scripts and environment specifications are available in the associated code repository, allowing for exact replication of the computational environment.

We recommend running the code examples within an isolated virtual environment, which can be created either with `conda` or the built-in `venv` module, to ensure dependency consistency across platforms. Detailed installation instructions are available from the official documentation for each tool:

- **conda:** https://docs.anaconda.com/anaconda/install/
- **venv:** https://docs.python.org/3/tutorial/venv.html

### 2.3 Data

The current protocol uses the FCON1000 dataset containing healthy participants as a reference sample, which is freely available at the [FCON1000 website](https://fcon_1000.projects.nitrc.org/indi/fcp_abcd/abcd.html). This dataset can be downloaded from https://fcon_1000.projects.nitrc.org/. Ethical approval for this study was provided by the institutional review boards for the study sites contributing data, and all participants provided informed written consent. For more details, please refer to the primary study paper [37] and website.

We will use cortical thickness imaging-derived phenotypes (IDPs) from several different datasets, which we list below. The datasets are all pre-processed and included in this repository. All preprocessing was done using [Freesurfer](https://surfer.nmr.mgh.harvard.edu/) with the `recon-all` command. We extracted the mean cortical thickness values following the Destrieux parcellation [38]. In addition, for the main protocol we will use two clinical datasets from the [OpenNeuro](https://openneuro.org/) portal.

1. ds003568: Contains data from 49 subjects (31F, 18M), aged between 12 and 19. Clinical labels are Healthy Control (HC, 20) and Major Depressive Disorder (MDD, 29) and individuals with comorbid or other disorders were excluded. For full details, such as ethical approvals and inclusion/exclusion criteria, please see [39].
2. ds003653: Contains data from 73 subjects (56F, 17M), aged between 19 and 44. Clinical labels are HC (39) and MDD (33) and individuals with comorbid or other disorders were excluded. For full details, including ethical approvals and inclusion/exclusion criteria, please see [40].

In addition, we use three additional and non-overlapping datasets to demonstrate the federated learning workflow:

1. ds003469: Contains data from 81 healthy participants (50F, 31M), aged between 18 and 39. For full details, see: doi:10.18112/openneuro.ds003469.v1.0.0
2. ds003826: Contains data from 136 healthy participants (87F, 49M) aged between 18 and 35. For full details, see: [41]
3. ds000222: Contains data from 79 healthy subjects (41F, 38M) aged between 21 and 73. For full details, see [42]

For the longitudinal workflow, we use an additional clinical dataset containing 98 individuals with first episode psychosis (38F, 60M, aged 18–46)) and 67 healthy controls (42F, 25M, aged 18-54). Each individual was scanned on two timepoints approximately one year apart. Full experimental details are provided in [11], but briefly individuals with psychosis were experiencing their first episode and individuals and comorbid conditions were excluded. Individuals with psychosis were treated by their consulting physician. All subjects provided written informed consent.

To illustrate the core workflow, in this protocol we show the results from one representative feature from the Freesurfer Destrieux atlas (*lh_G_and_S_frontomargin*). This feature was chosen as a typical example with a moderate age effect, heteroskedasticity, and variance across sites. Because convergence behavior is generally consistent across features with comparable sample sizes and data characteristics, diagnostics for this representative case serve as a practical indicator of overall model stability.

## 3 Procedure

### Installation and verification Timing: ~1-2 min

1. Install PCNtoolkit in the virtual environment created following the instructions in Section 2.2:

`!pip install pcntoolkit`
2. Verify the installation by printing the version string: 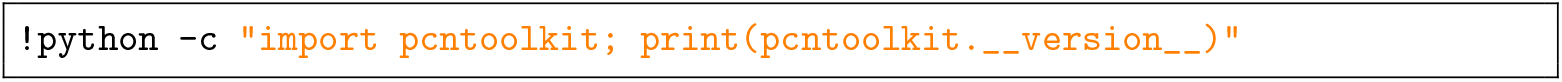

### Data selection and loading TIMING: ~1 min

3. Load the reference and clinical datasets into pandas DataFrames. For the estimation of the normative model, we will use the FCON1000 dataset and then apply these models to two clinical datasest from OpenfMRI. See 2.3 for details. To achieve this, execute: 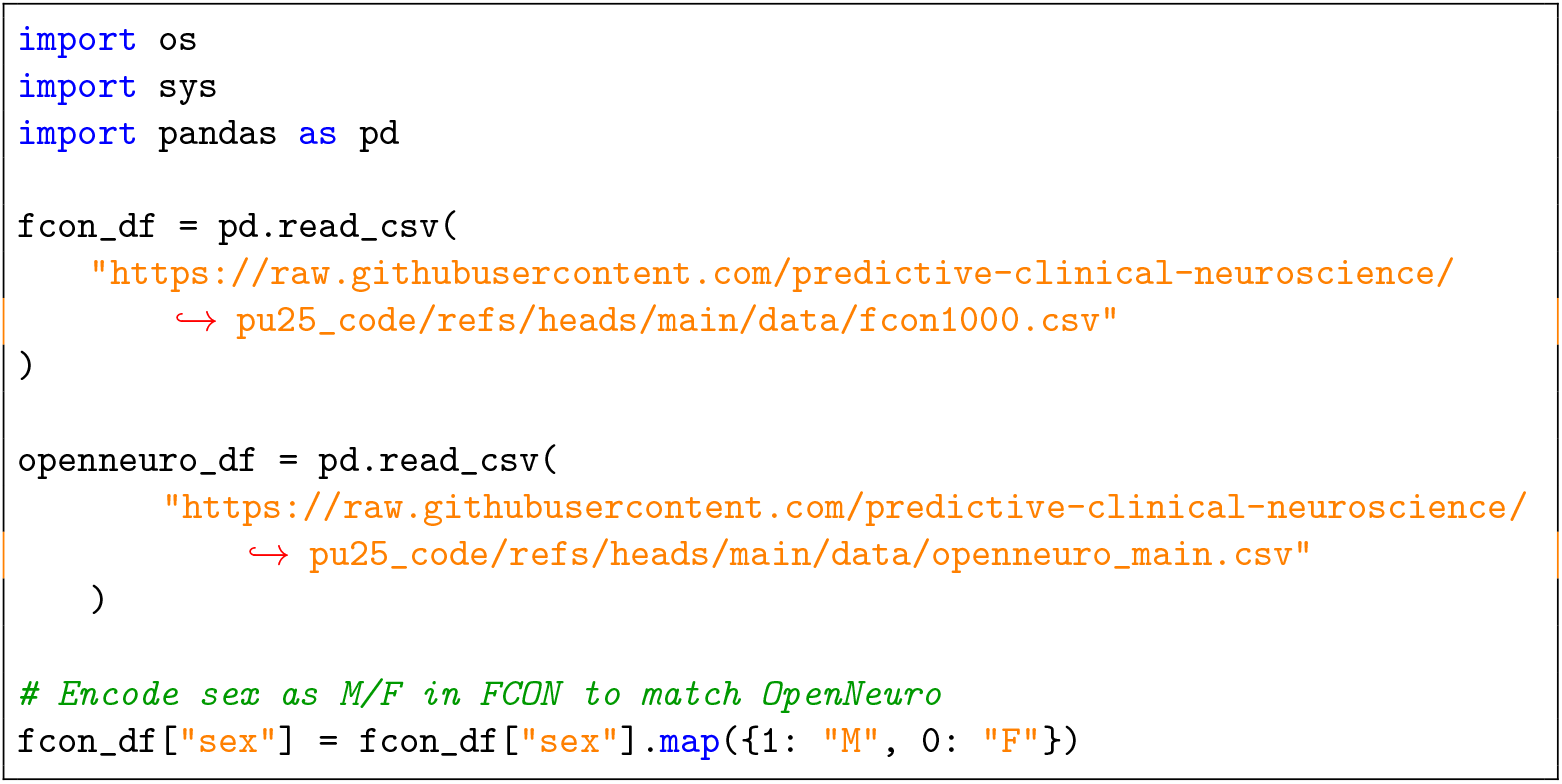
4. Define the subject identifier, covariates, batch effects, and response variables for the analysis (see Section 1.5 for guidance on these choices): 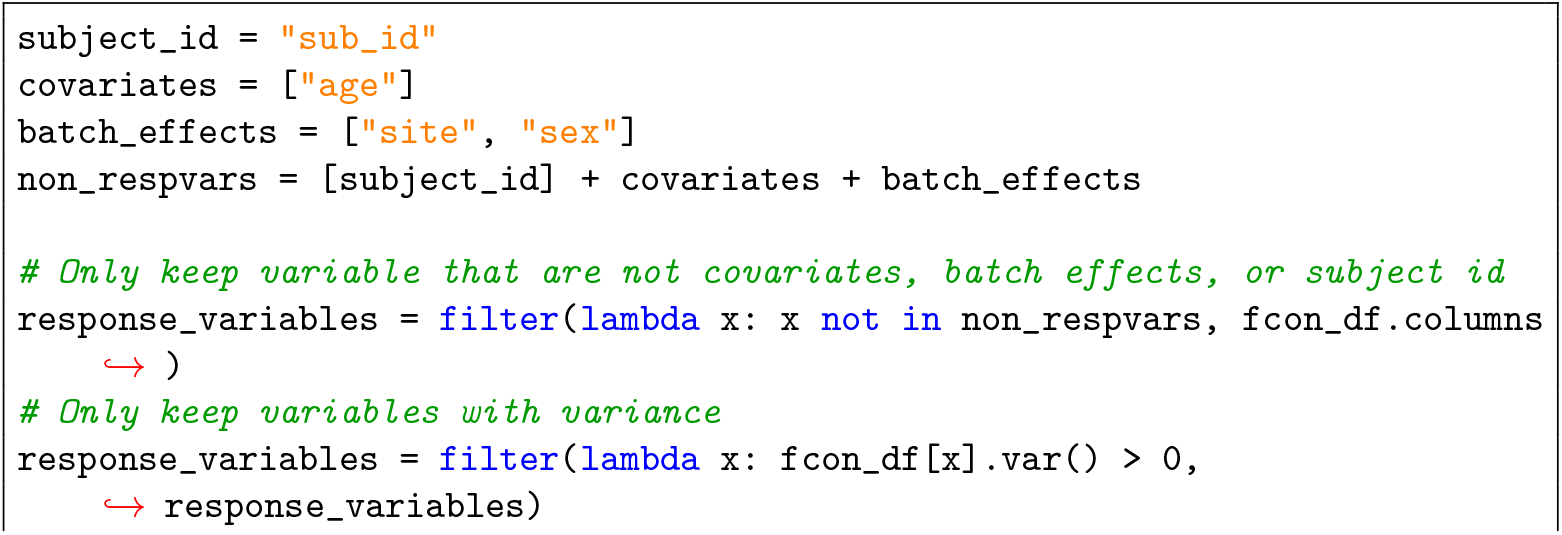
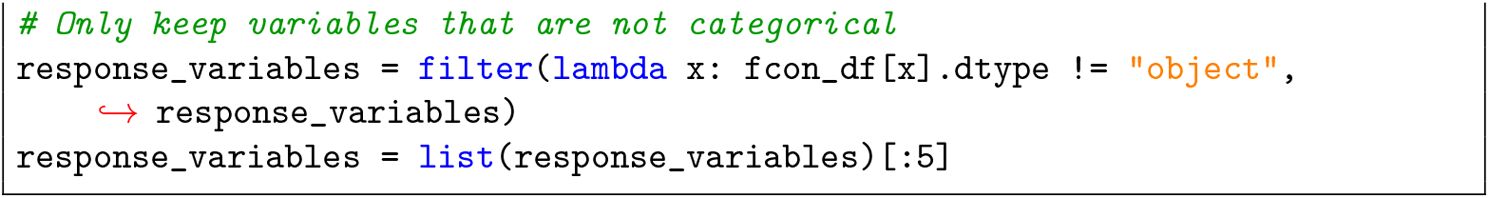

### Data preparation TIMING ~1 min

5. Create a `NormData` object from the reference dataframe, specifying outlier removal and missing-value handling (see Section 1.5 for guidance on `z_threshold` selection).

#### CRITICAL STEP

Set `z_threshold` appropriately for your data. Overly aggressive values (e.g. 3) can truncate the empirical distribution; overly liberal values (e.g. 10) retain more data but may permit extreme values. See 1.5 for considerations about the threshold level.

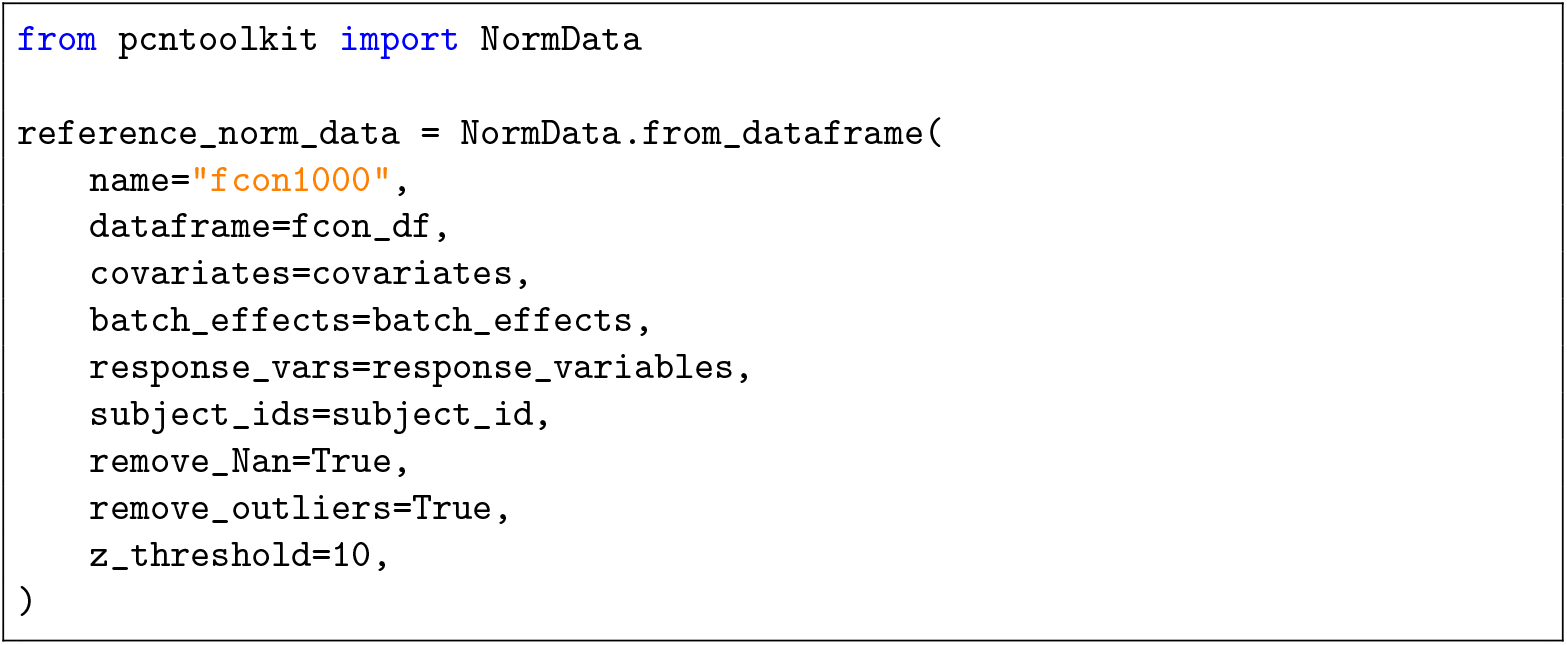

### Combining data sources TIMING ~1 min

6. Create separate `NormData` objects for each clincial dataset and split by clinical group: 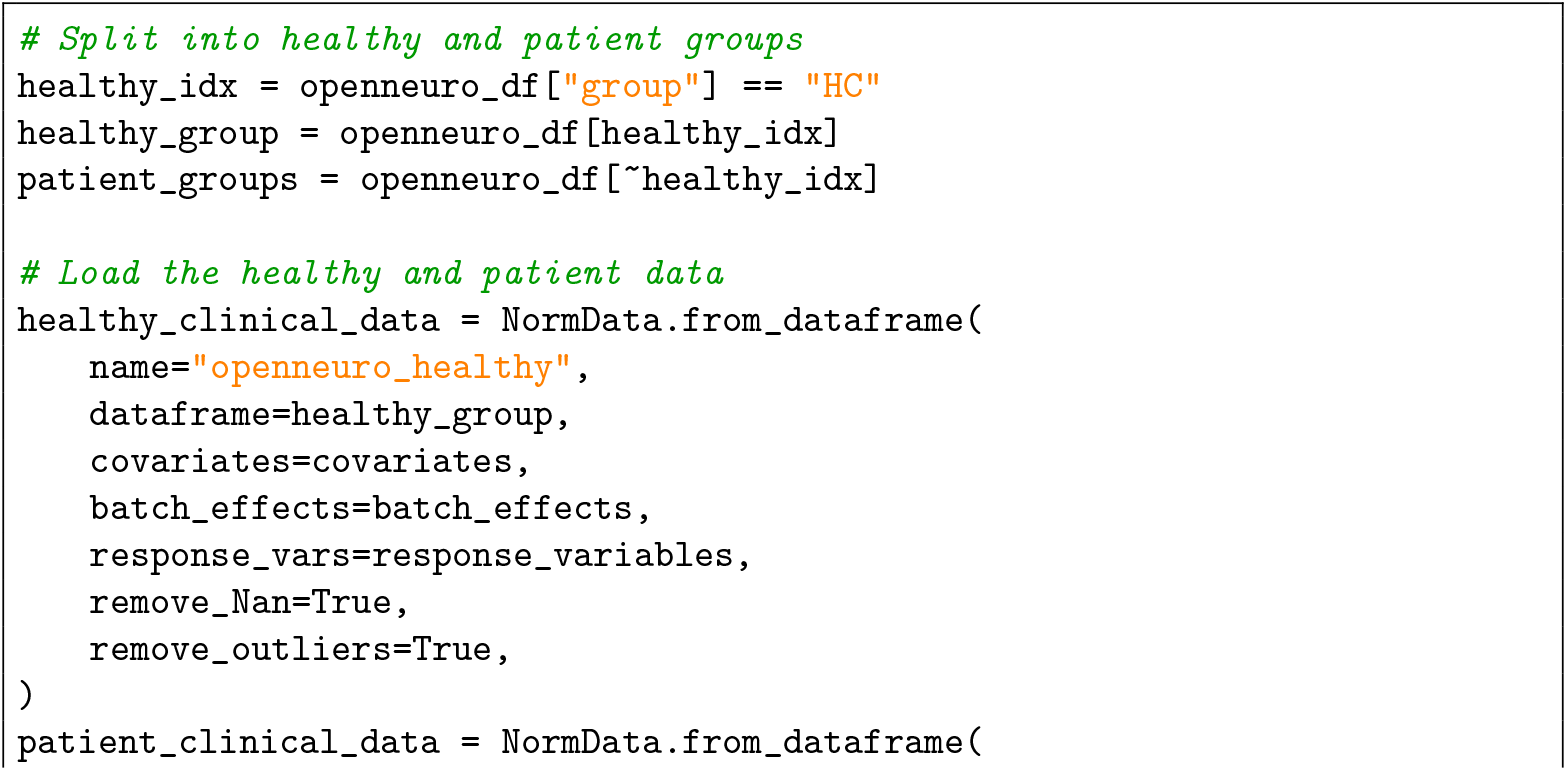
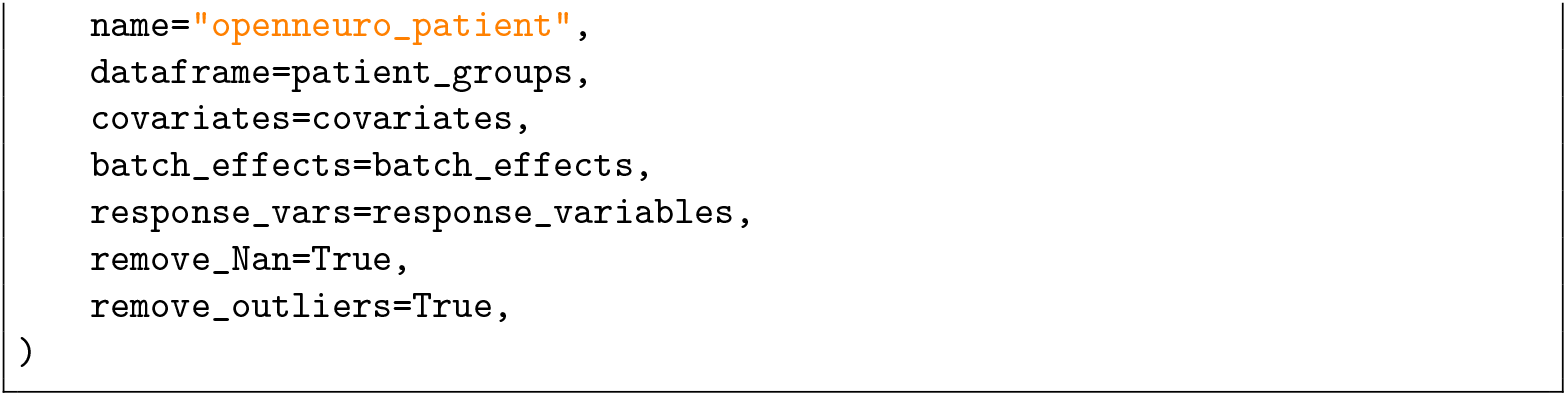
7. Ensure batch-effect compatibility between datasets and merge.

`reference_norm_data.make_compatible(healthy_clinical_data)`

`norm_train = reference_norm_data.merge(healthy_clinical_data)`
8. Verify schema compatibility of the clinical (patient) data against the merged reference set: 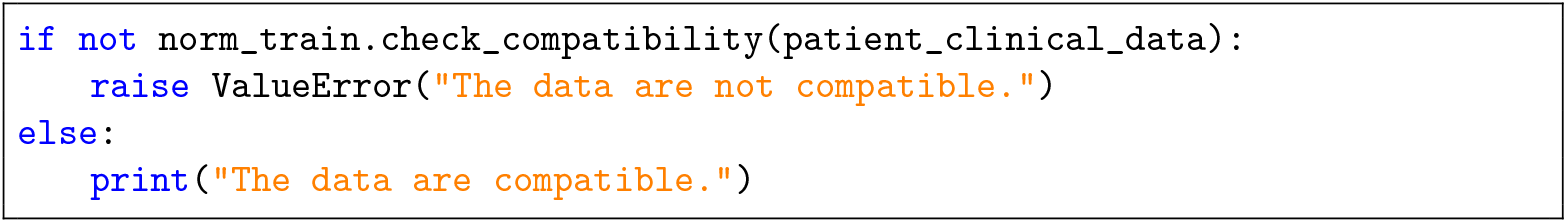

### Train–test split TIMING ~1 min

9. Split the merged reference data into training and test sets using the `NormData.train_test_split` method, which internally calls the `scikit-learn` function of the same name. See Section 1.5 for guidance on split ratios and stratification. 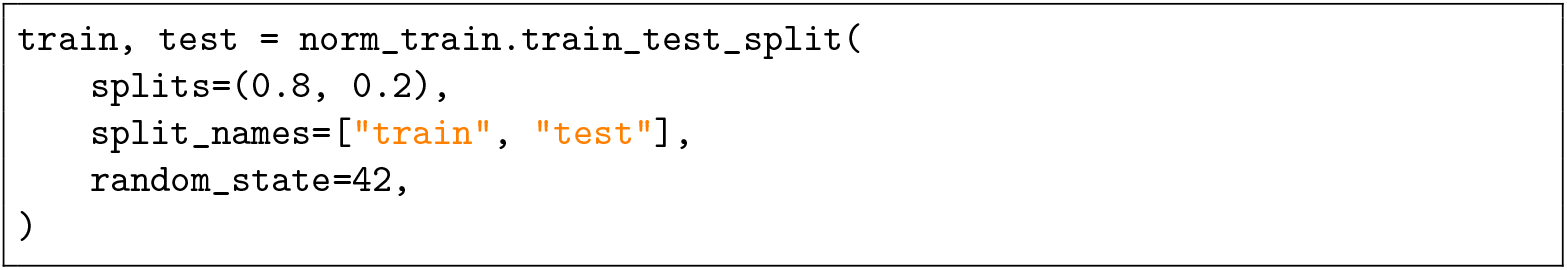

### Model configuration and fitting TIMING ~5–15 min

10. Configure and instantiate the normative model. Select one of the following options:
  A. *Default HBR configuration.* Use this for an initial exploration with HBR with default parameters. In the model comparison workflow, we compare the suitability of thsi mode to the more complex model below. 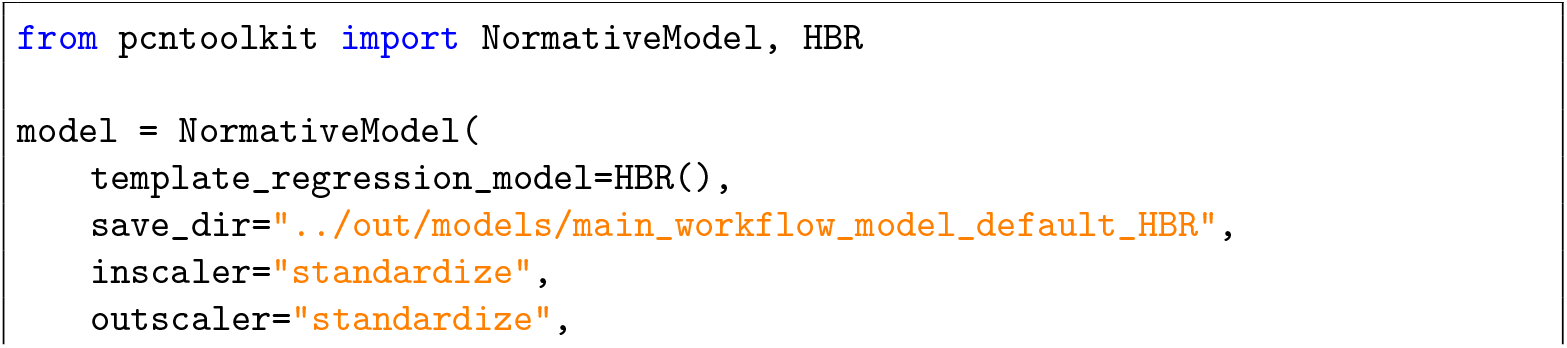
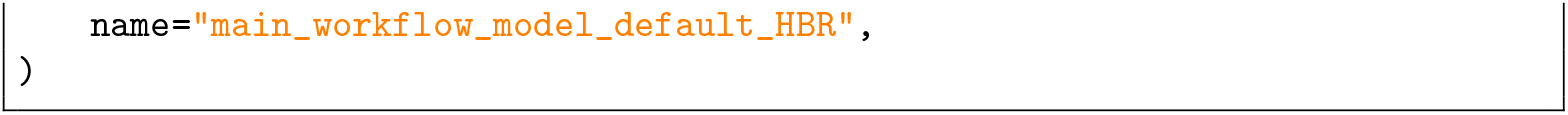
  B. *Custom HBR with specified likelihood and priors.* Define a hierarchical Normal model with B-spline basis functions, random intercepts for site and sex, and heteroskedastic variance (see Section 1.5.3 for guidance on likelihood and prior choices).

In this workflow, we use the Normal likelihood, which assumes the residuals follow a Gaussian distribution after preprocessing and model fitting. This is appropriate for cortical thickness data, where deviations are approximately symmetric and unbounded. Both *µ* and *σ* are modeled hierarchically as functions of age (covariate) with random intercepts for site and sex, allowing age effects to vary smoothly while accounting for systematic offsets across acquisition sites and sexes (batch effects).

To achieve this, we define a hierarchical Gaussian model where each observation *y*_*n*_ is generated according to equations 1–6, that is, according to the generative process implemented in the PCNtoolkit’s HBR class. For full details please see [9], but briefly, each response variable is modeled with a Gaussian likelihood whose mean (*µ*_*n*_) and standard deviation (*σ*_*n*_) depend on a basis expansion of the covariates (*ϕ*(*x*_*n*_)) (equations 2 and 3). A softplus mapping function is employed to ensure the standard deviation remains positive (*σ*^+^ = ln(1 + exp(*σ/*3)) ∗ 3). Finally, equation 4 describes a reparameterisation whereby batch effects are incorporated by combining an overall mean 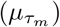 with the group standard deviation 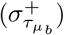 sampled from a positive distribution with an offset for each level of the batch effect 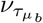. As outlined in [43], this parameterisation improves sampling efficiency.

Concretely, define the prior structure for the mean (*µ*) and scale (*σ*) parameters:

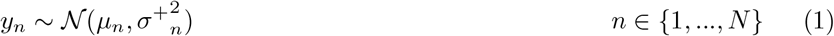

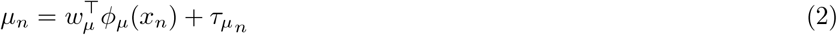

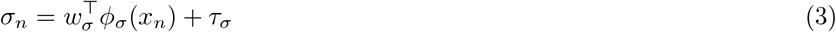

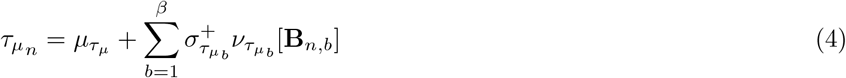

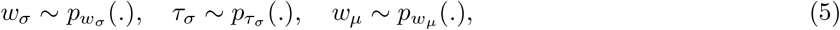

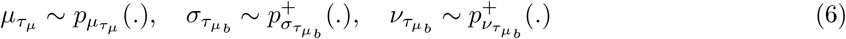

where:

- *N* is the number of observations.
- *w*_*µ*_ and 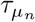 are respectively regression coefficients and standard deviations for the linear model for the mean.
- *w*_*σ*_ and *τ*_*σ*_ are respectively regression coefficients and standard deviations for the linear model for the variance.
- *β* is the number of batch effect dimensions, i.e., for batch effects (site, sex), *β* = 2.
- **B** denotes the *N* × *β* matrix of batch effect indices.
- 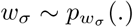 is a Gaussian prior over 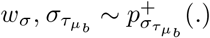 is a softplus-transformed Gaussian prior for 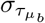, and similarly for the other variables. We omit the parameters of these distributions for brevity.

To configure this in PCNtoolkit, run the following code:

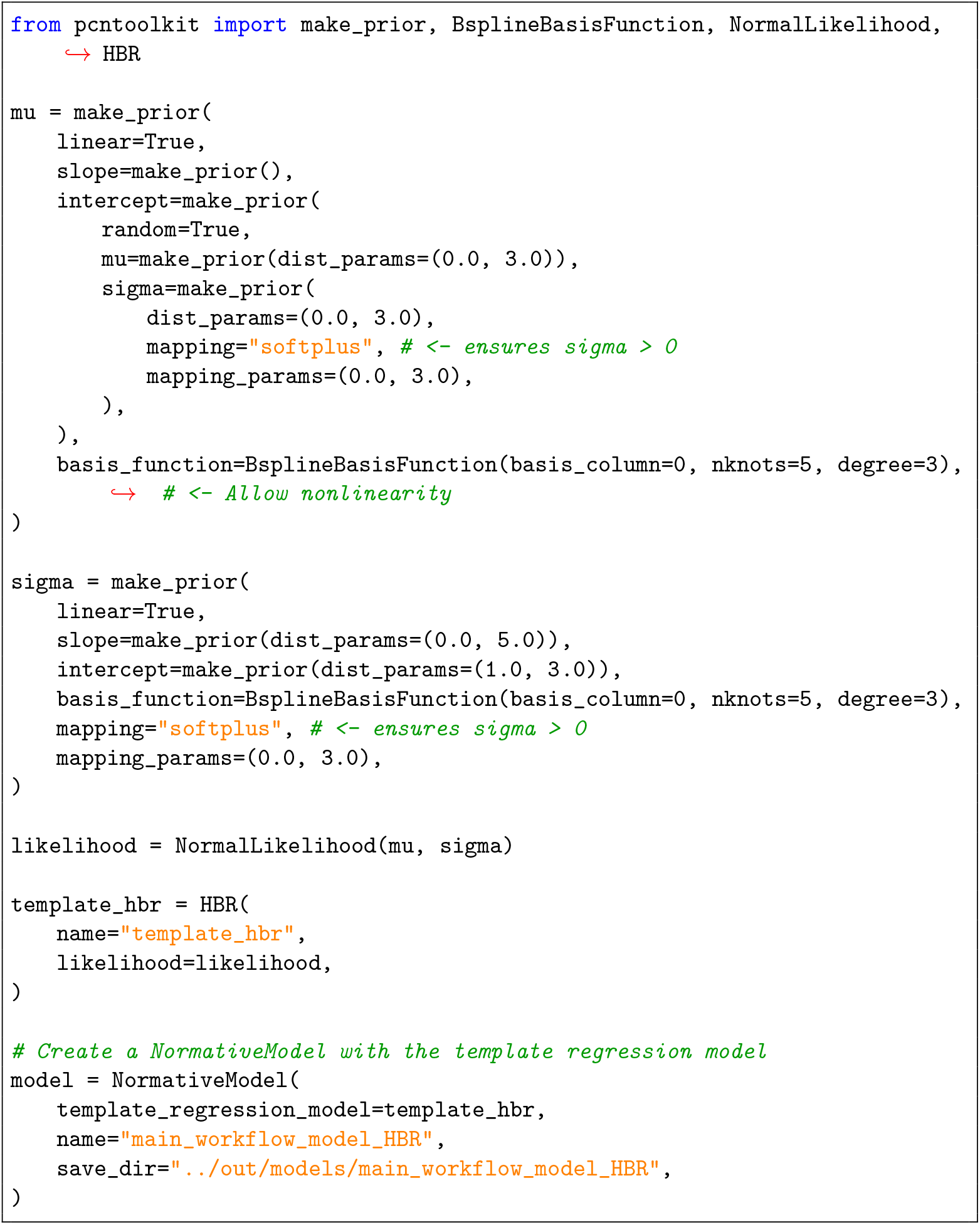

11. Fit the model on the training set and generate predictions for the test set. Select one of the following options:
  A. **Local fitting.** For small-to-moderate datasets on a single machine:

`model.fit_predict(train, test)`
  B. **Parallel fitting on a compute cluster.** For large datasets with access to a Slurm or Torque workload manager. 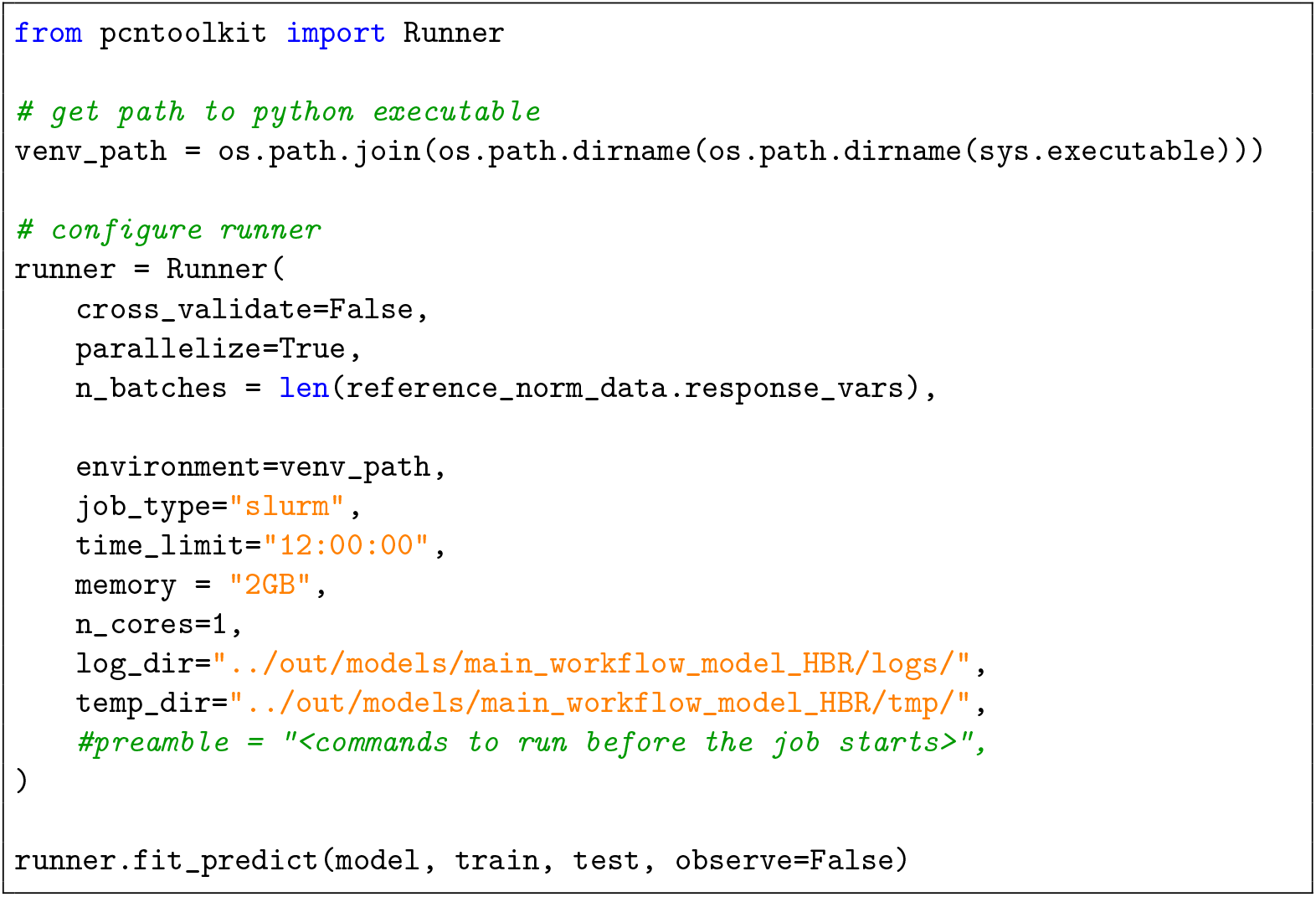

#### PAUSE POINT

The fitted model and all outputs are saved to disk in the directory specified by `save_dir` (see Section 4 for the output directory structure). The workflow can be resumed from the saved model at any later time.

### Model evaluation TIMING ~5–10 min

12. Inspect convergence diagnostics for all fitted features. This is necessary for HBR (but not BLR). Load posterior summaries from the per-feature `idata.nc` files and verify that all parameters meet the following standard MCMC diagnostics. More specifically, we require all parameters to have potential scale reduction factors 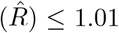, bulk and tail effective sample sizes (ESS) ≥ 1000, and Monte Carlo standard errors (MCSE) below 1% of the corresponding posterior standard deviation. Visually inspect the chains to confirm stationary and adequate mixing across draws. ^2^See section 4 for examples.

#### CRITICAL STEP

Models that fail these convergence criteria should be regarded with caution.

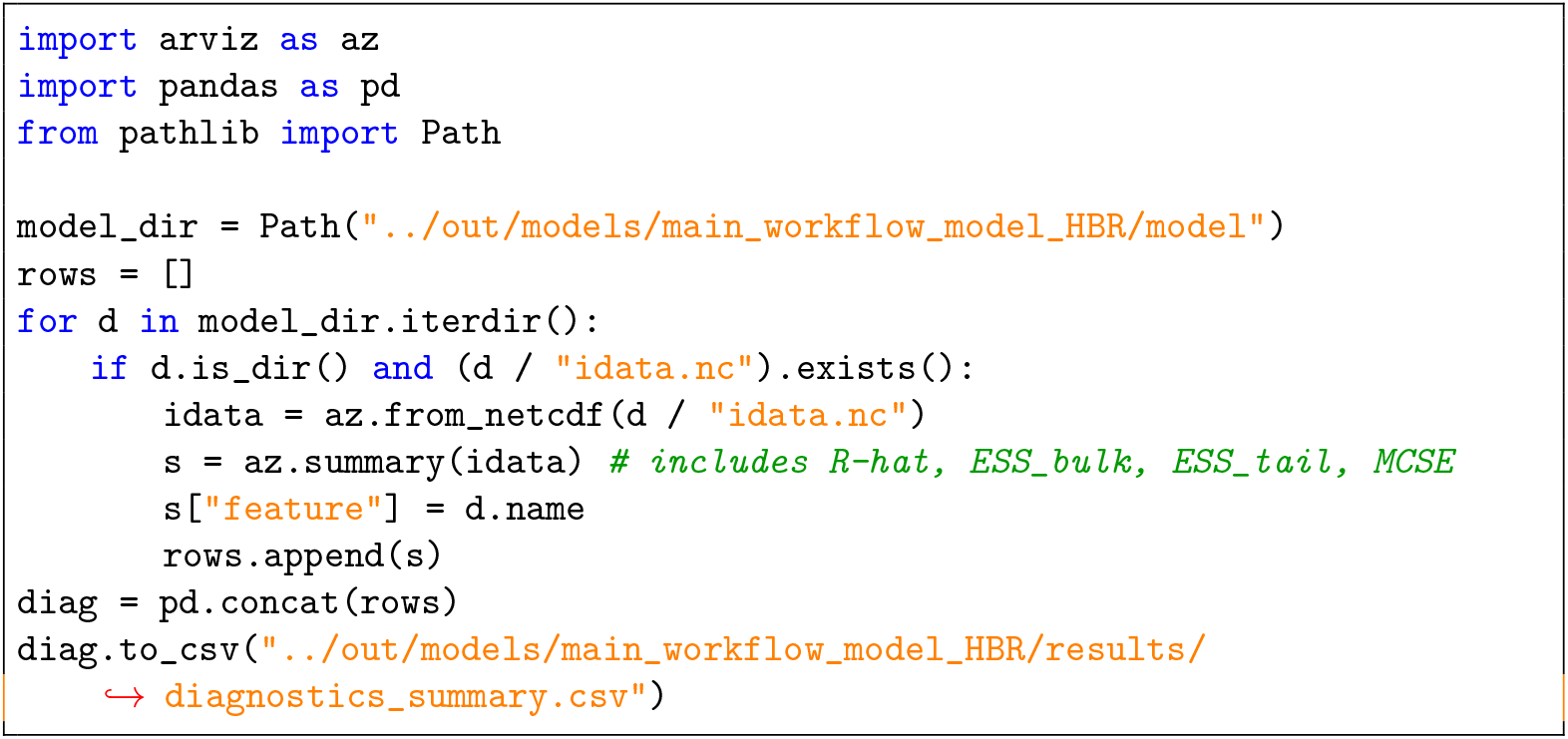

13. Load and review quantitative evaluation metrics (see Anticipated Results for interpretation): 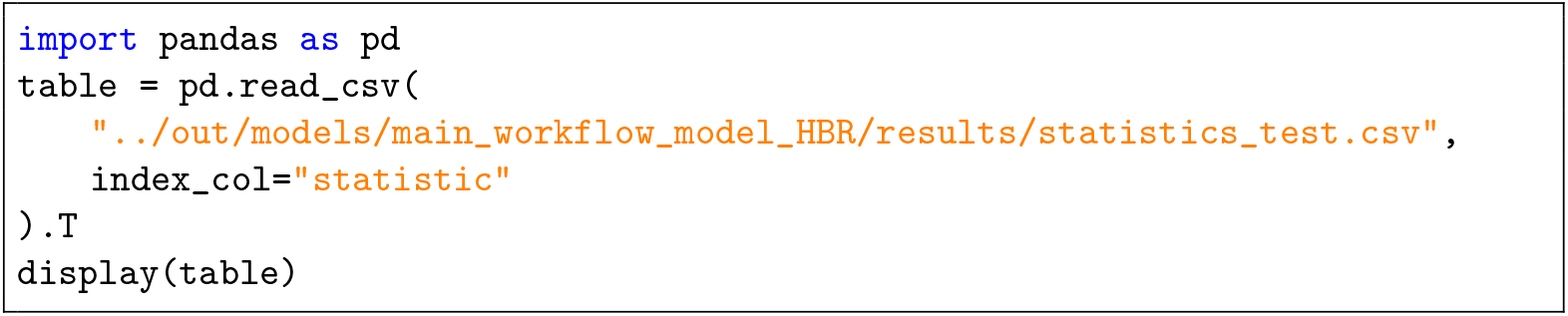
14. Generate centile plots and QQ plots for visual calibration assessment (see section 4 for examples and interpretation): 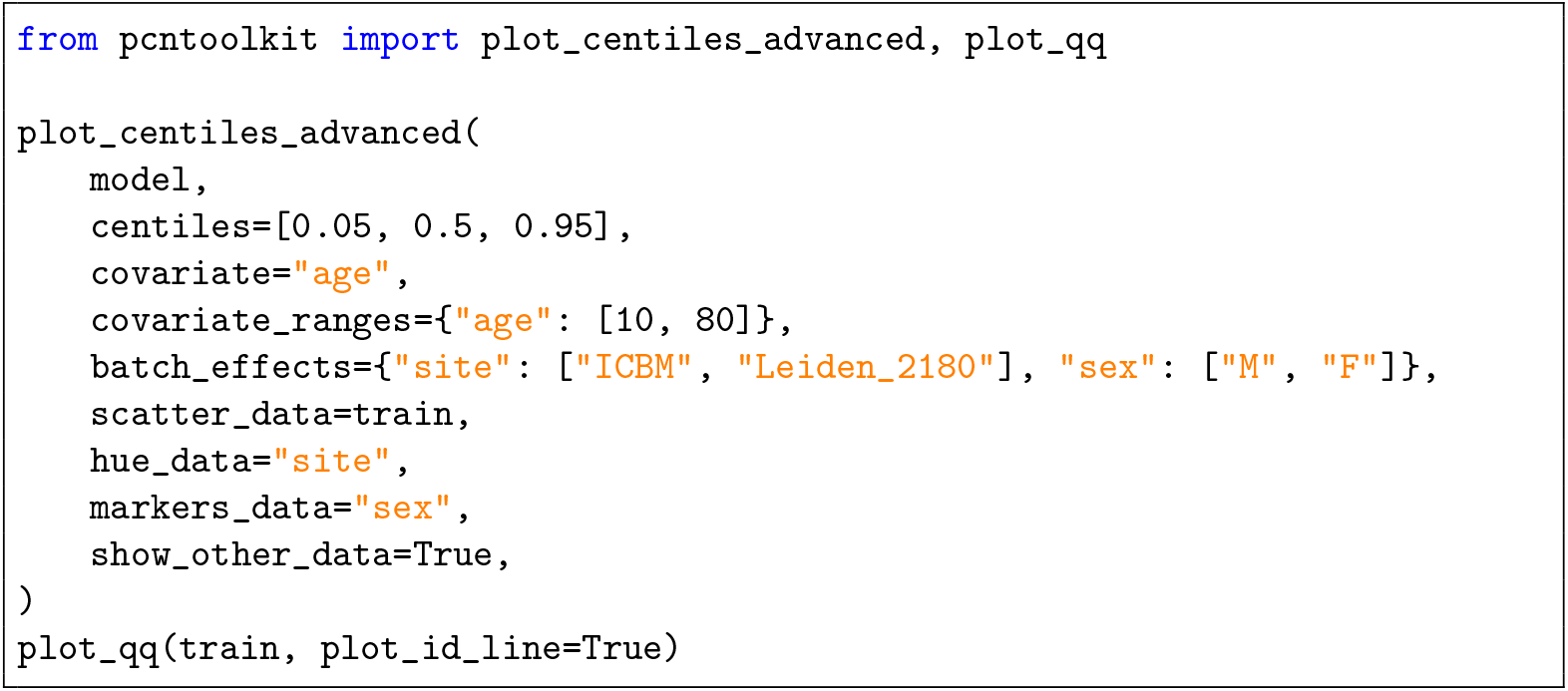

#### PAUSE POINT

At this stage of the protocol, the basic workflow is complete. With convergence and calibration verified, the model is ready for downstream applications such as longitudinal analyses, data harmonization, and federated normative modeling on decentralized data. The additional workflows provided in the sections below explain several of these analyses and should be executed as required, not sequentially. For example, it is not necessary to first run the longitudinal normative modelling workflow in order to run the Federated Learning workflow.

### Optional secondary workflows

The secondary workflows that follow are simplified versions of the secondary workflow [notebooks](https://github.com/predictive-clinical-neuroscience/pu25_code/blob/main/notebooks/secondary_workflows) available as part of the associated repository, which is intended to highlight the underlying workflow. Rather than cutting and pasting the code below, we recommend cloning the associated repository, which provides additional flexibility and more extensive error handling.

### Optional Workflow: Federated Learning TIMING ~5–10 min

This workflow can be used to apply a pre-trained model to a new target dataset. Whilst we could use the pretrained model estimated in the main workflow, in this example, we instead choose a more realistic example. We pull a large scale pre-estimated model from our pre-estimated library (essentially an update to the cortical thickness models in [4]). The target dataset is from OpenNeuro and has the same cortical thickness measurements as the original dataset. Only the pre-trained model is required for this step. The original data are not accessed, preserving data privacy. For more extensive examples, please see the associated code repository, where we also provide an additional simplified workflow.

1. First, install the necessary packages and specify configuration variables. 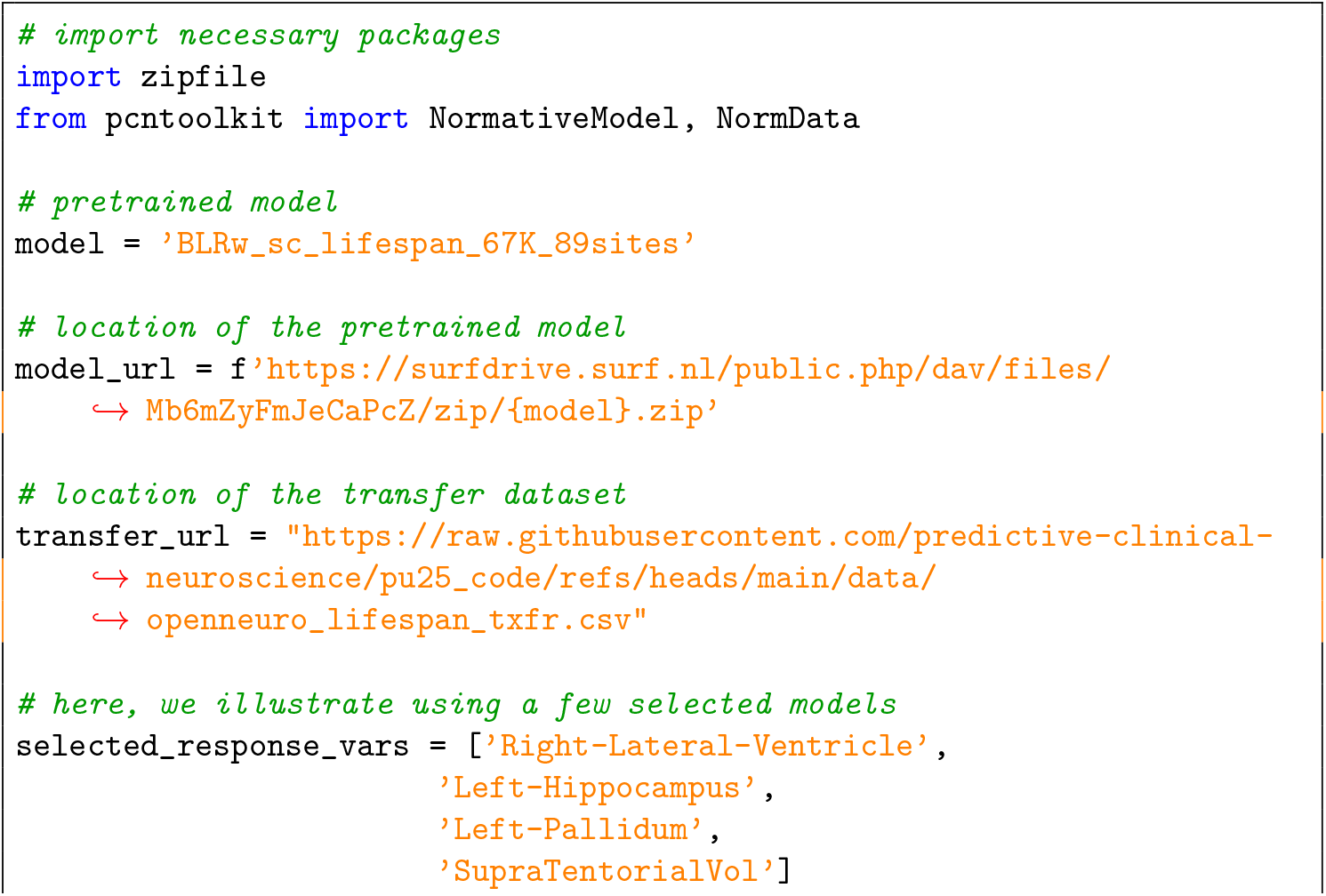
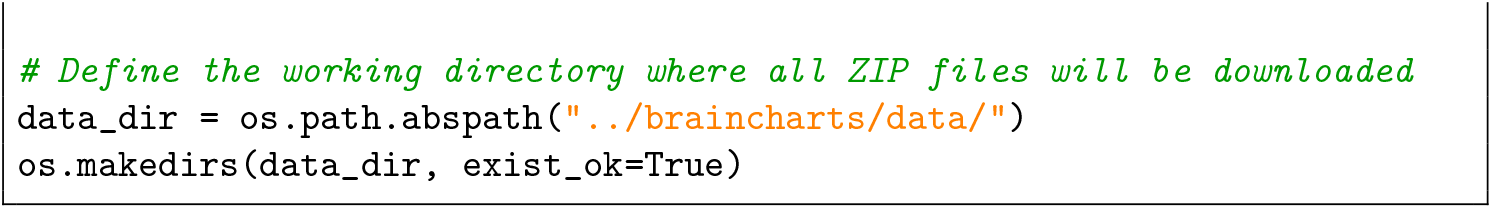
2. Next, download and unzip the pretrained normative models 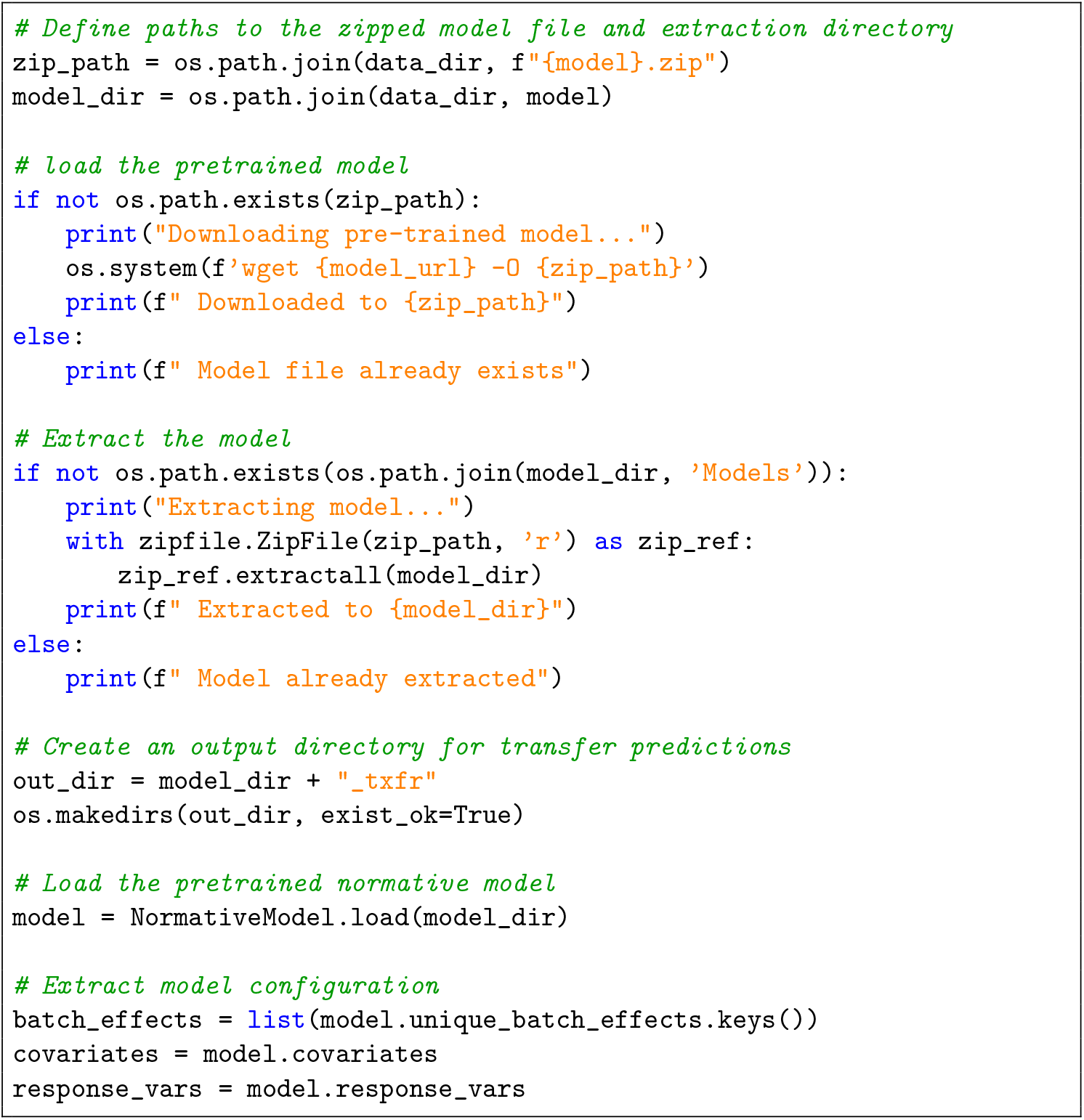
3. Load and prepare the transfer datasets (in this example case from OpenNeuro) 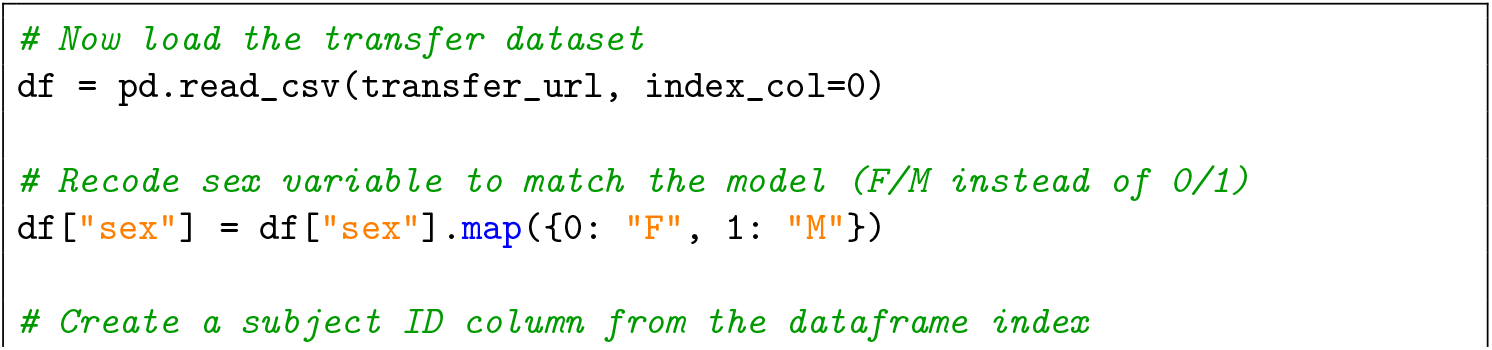
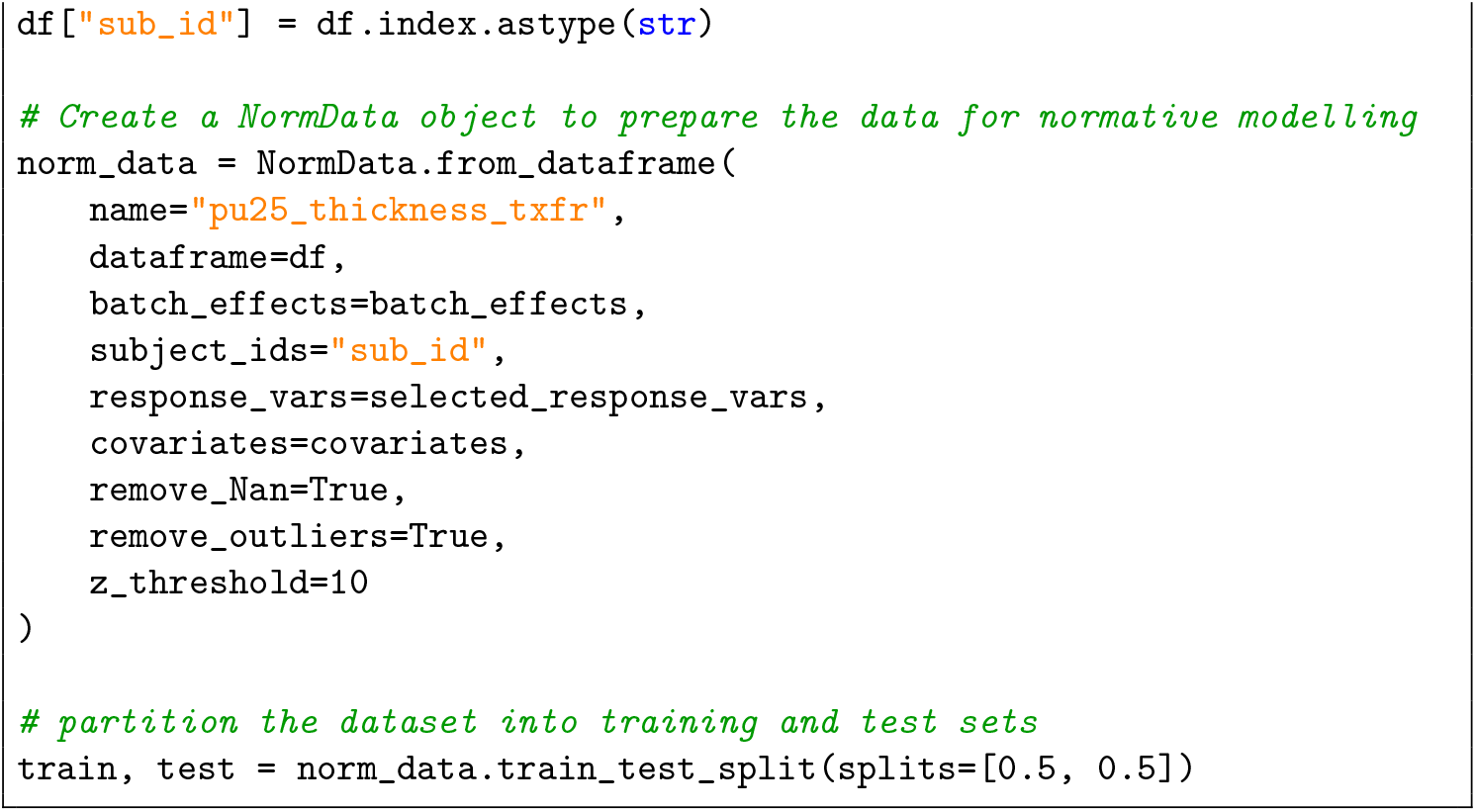
4. Finally, run the transfer prediction using PCNtoolkit routines 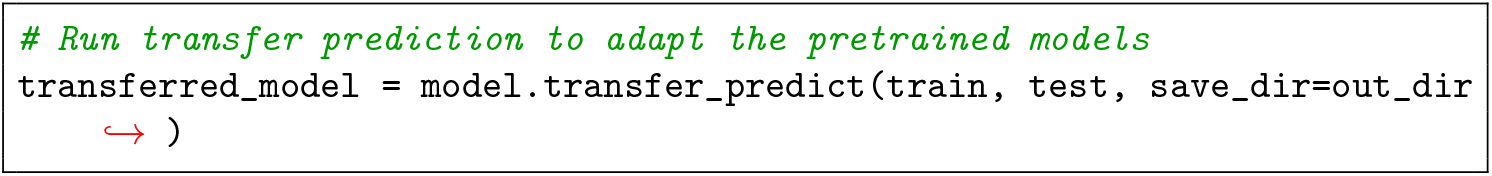

**PAUSE POINT:** At this point, the federated workflow outlined below is complete and the model metrics and centiles can be plotted (see steps 12 and 13 in the main procedure).

### Optional Workflow: Longitudinal Modeling TIMING ~15–20 min

This section demonstrates how to compute the z-diff score [11] to quantify longitudinal change in neuroimaging data relative to healthy ageing trajectories. We use a large-scale pre-trained BLR normative model [4, 13] from our pre-estimated library fitted on cortical thickness measures from the Destrieux parcellation across the lifespan (as is done in the federated learning workflow above). The target dataset consists of longitudinal structural MRI data from patients and healthy controls, with cortical thickness extracted using FreeSurfer, also documented in the original publication [11]. Rather than re-estimating the normative model on the new dataset, we leverage the pre-trained model directly by exploiting the fact that site-specific biases cancel out in the longitudinal difference, and compute z-diff scores that capture individual-level deviations from the expected healthy trajectory.

1. First, load the necessary packages and the longitudinal dataset 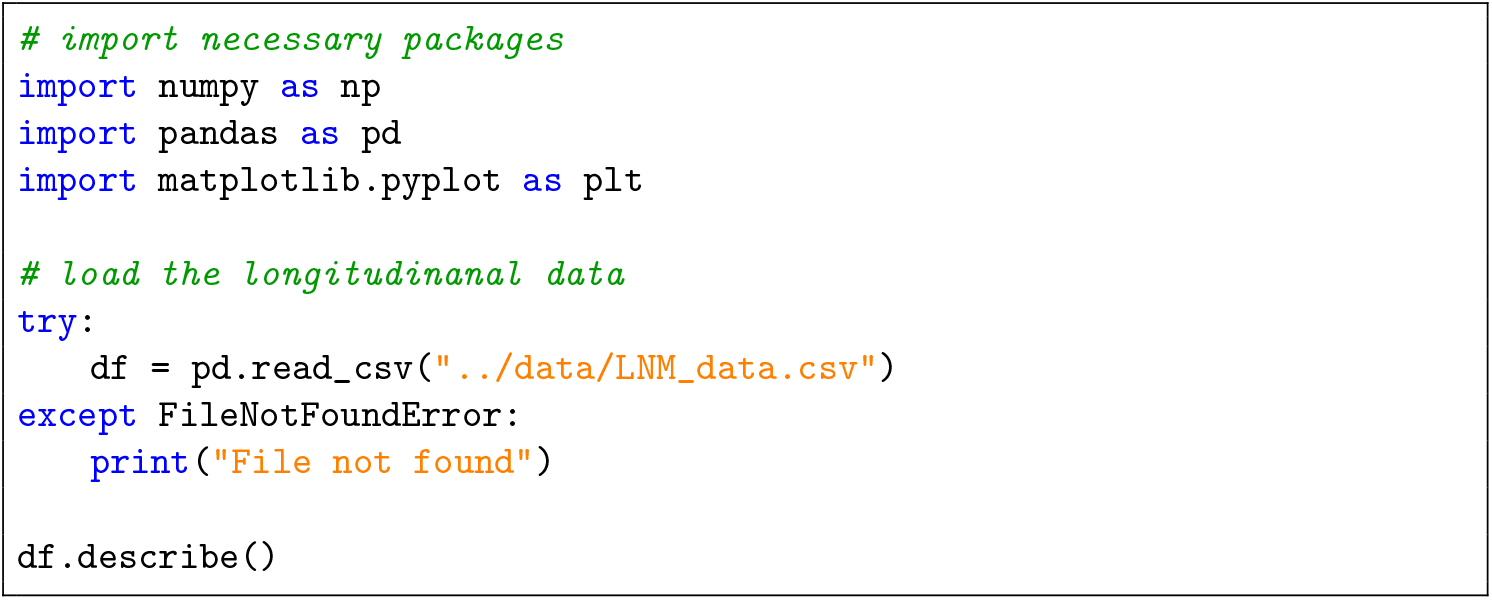
2. Next, download and unzip the pretrained normative models 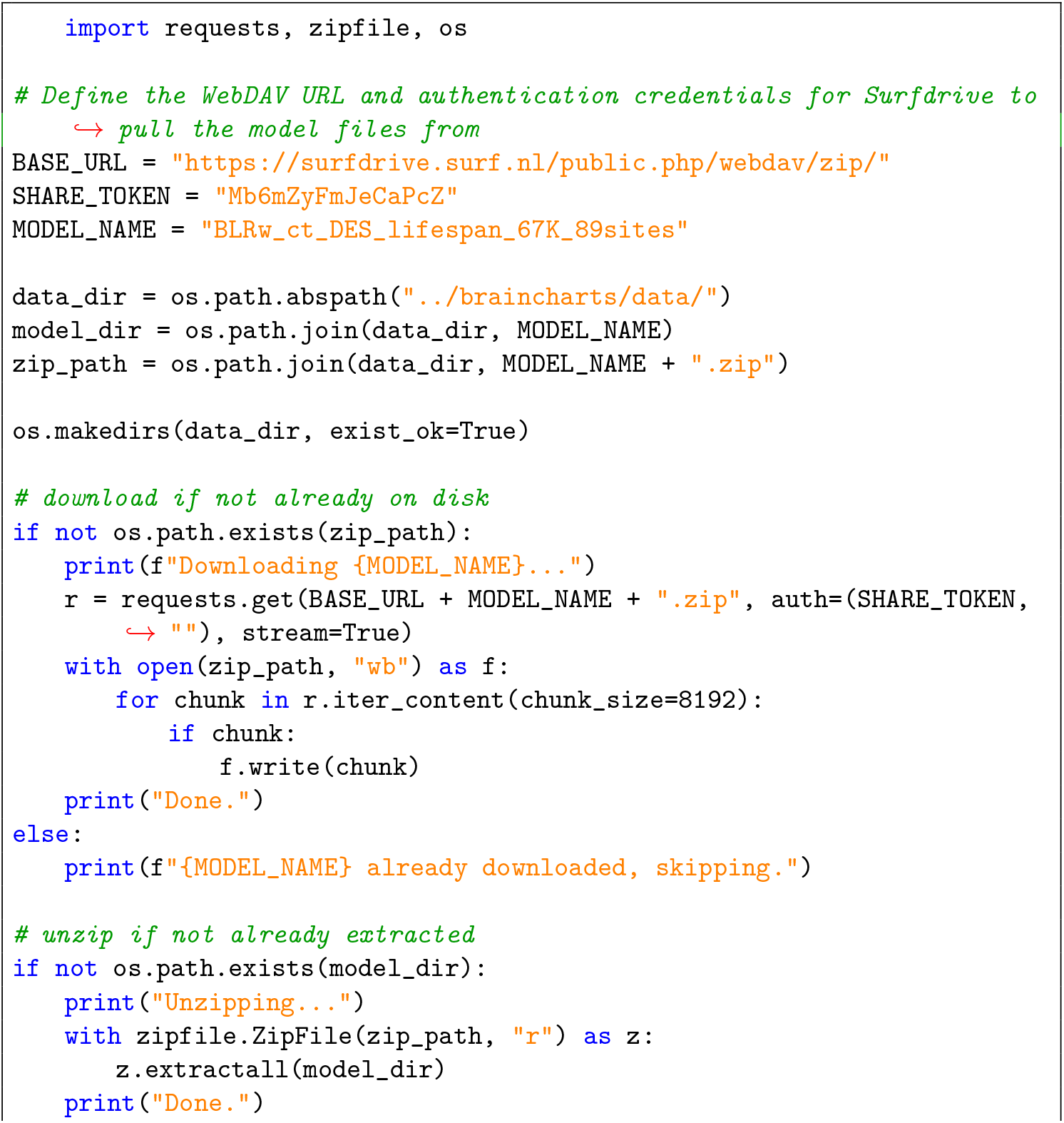
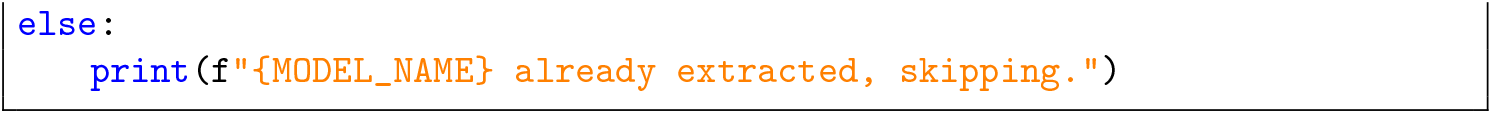
3. We load the pretrained model and define the covariates, batch effects, and response variables. 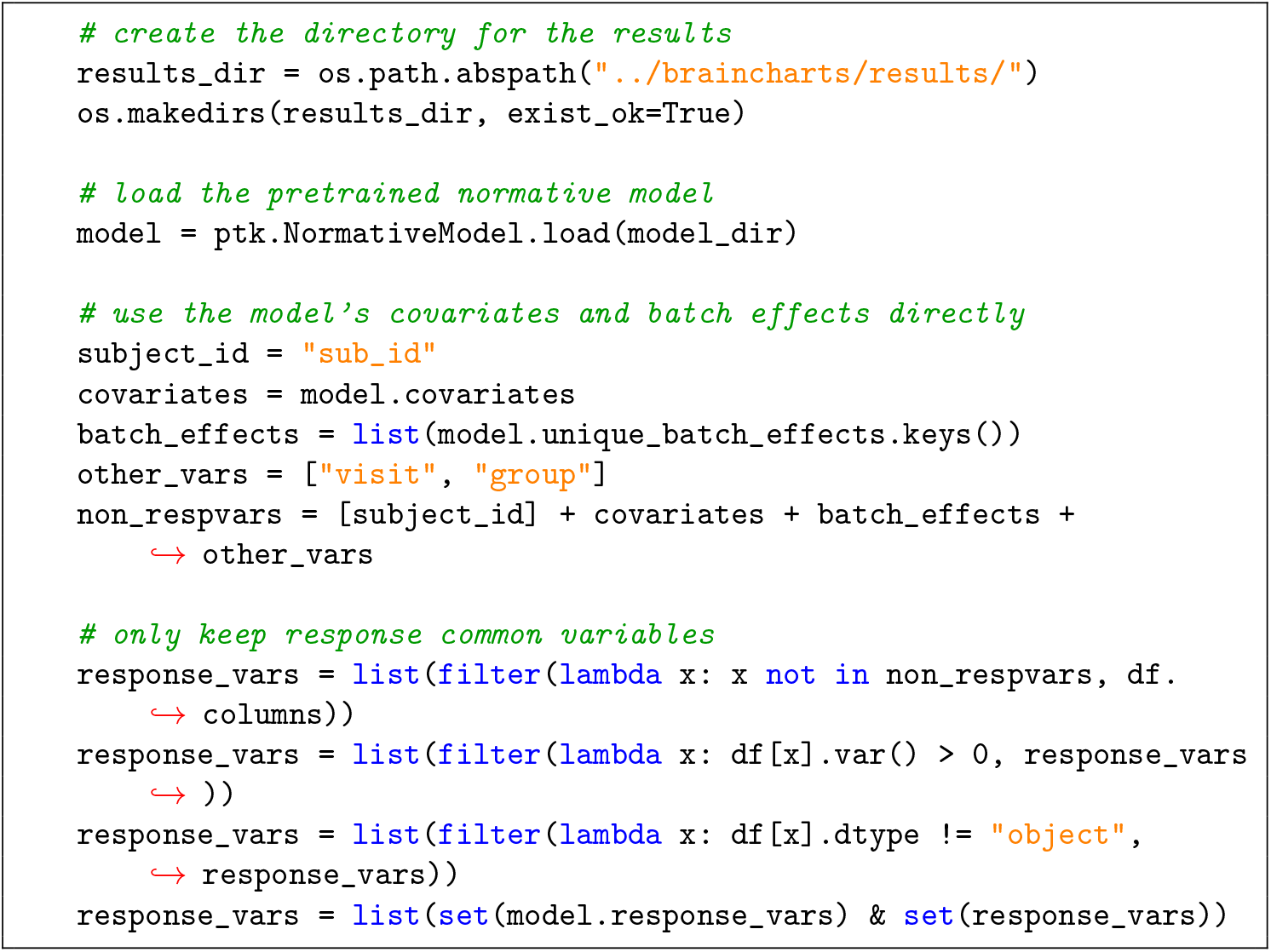
4. We split the data across groups and visits into separate `NormData` objects. Additionally, we also split the controls into an adaptation and transfer set – to estimate the site effect, and healthy between-visit variance, respectively. 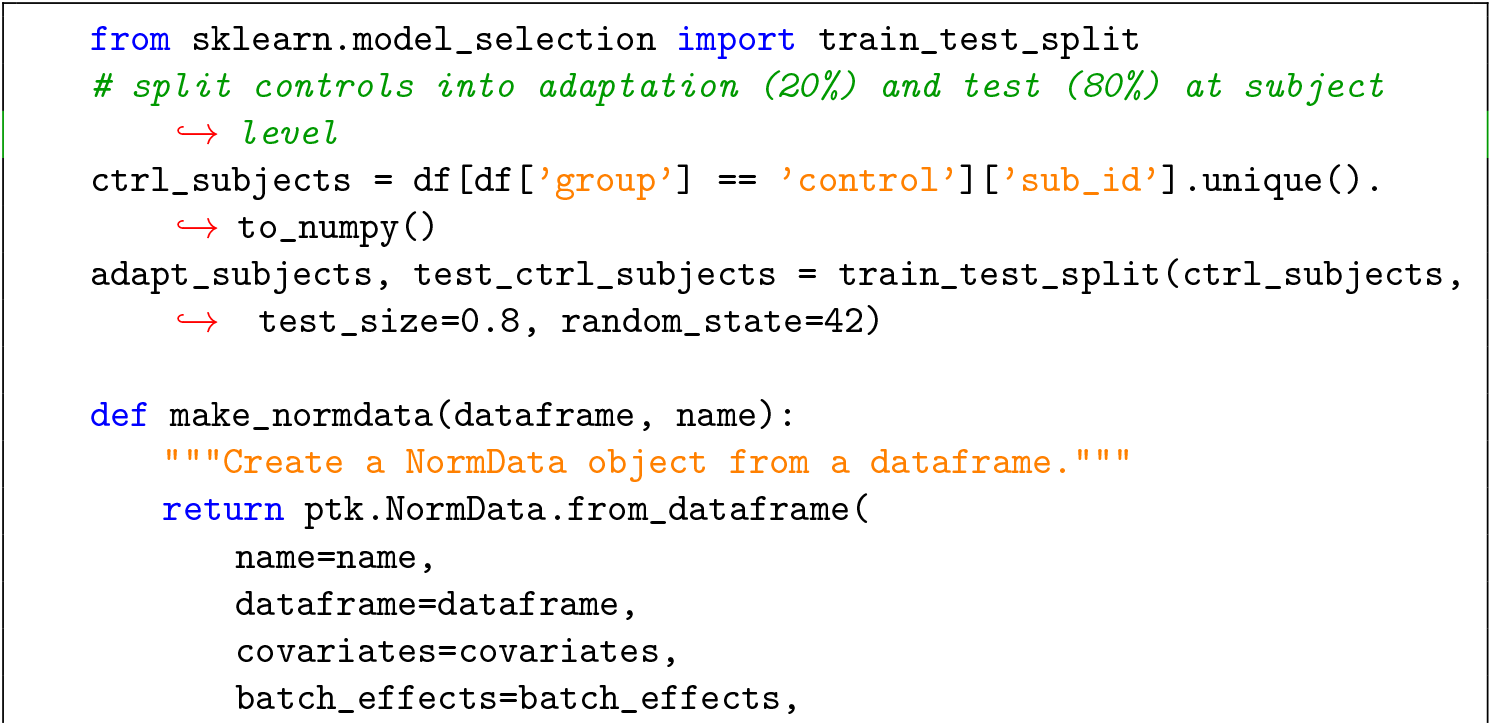
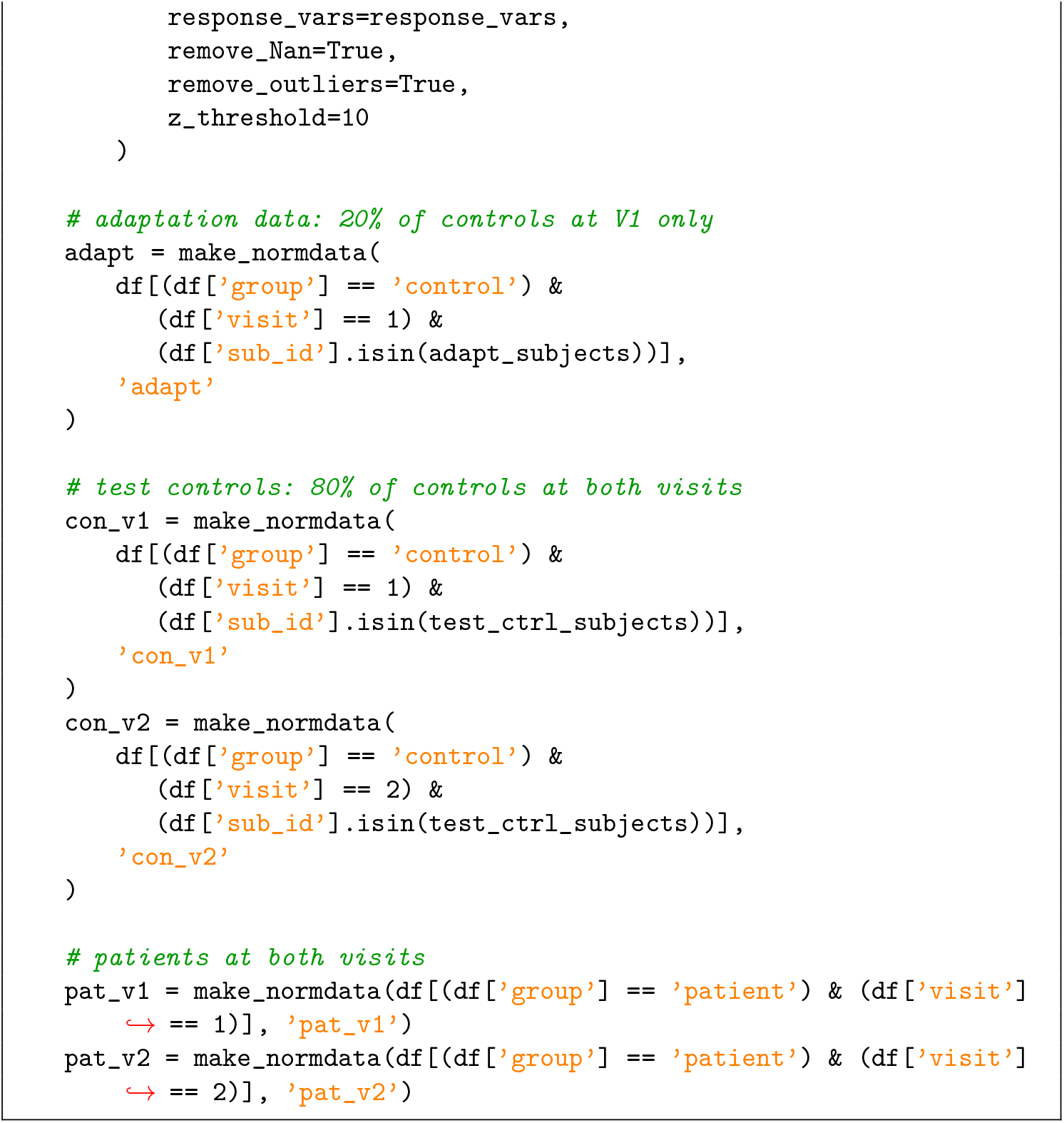
5. Transfer the pretrained model to the new site using the adaptation set and then use it to predict to both groups and visits. 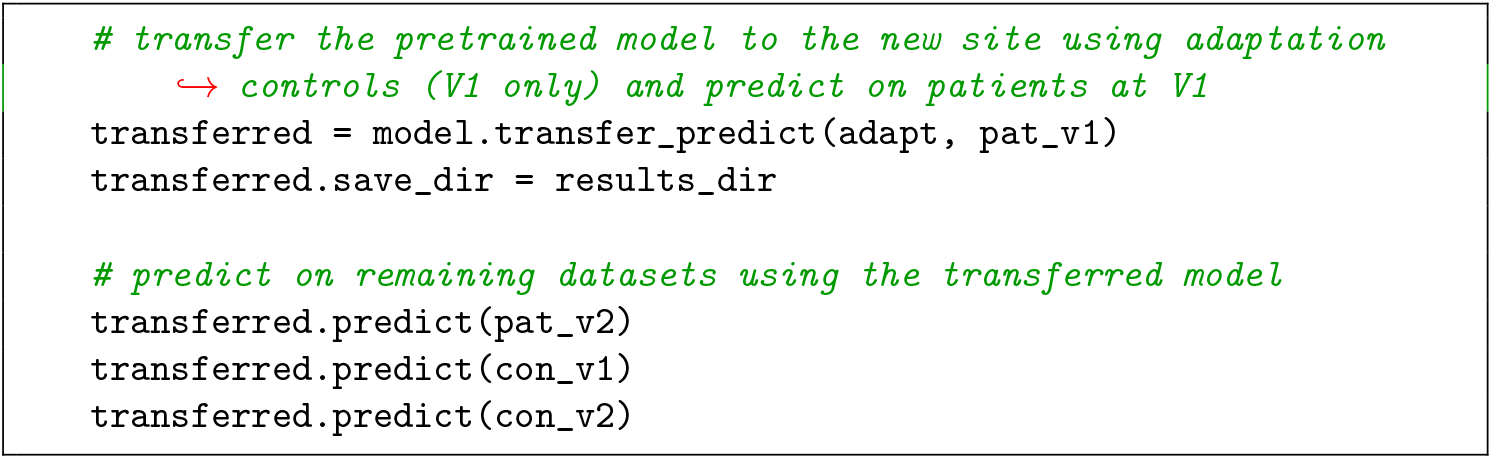
6. Transform predictions to warped space: The PCN toolkit returns predictions in the original (native) space rather than the warped space. However, the z-diff score must be computed in the warped space. We therefore apply the sinh-arcsinh warp to both the observed values *y* and the predicted values *ŷ* to recover the warped residuals *φ*(*y*) − *φ*(*ŷ*), which form the numerator of the z-diff equation. 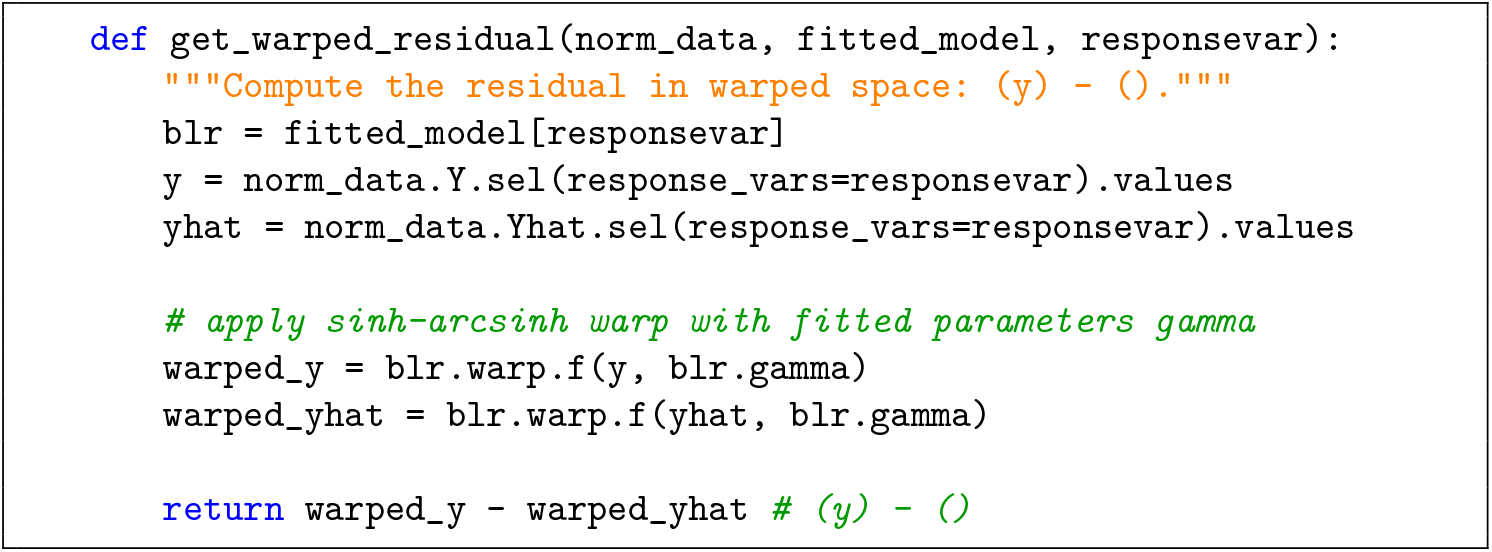
7. Z-diff score computation: The z-diff score quantifies the longitudinal change in a brain measure relative to what would be expected based on healthy ageing, normalized by the uncertainty in that prediction (see 1.5 for explanation. 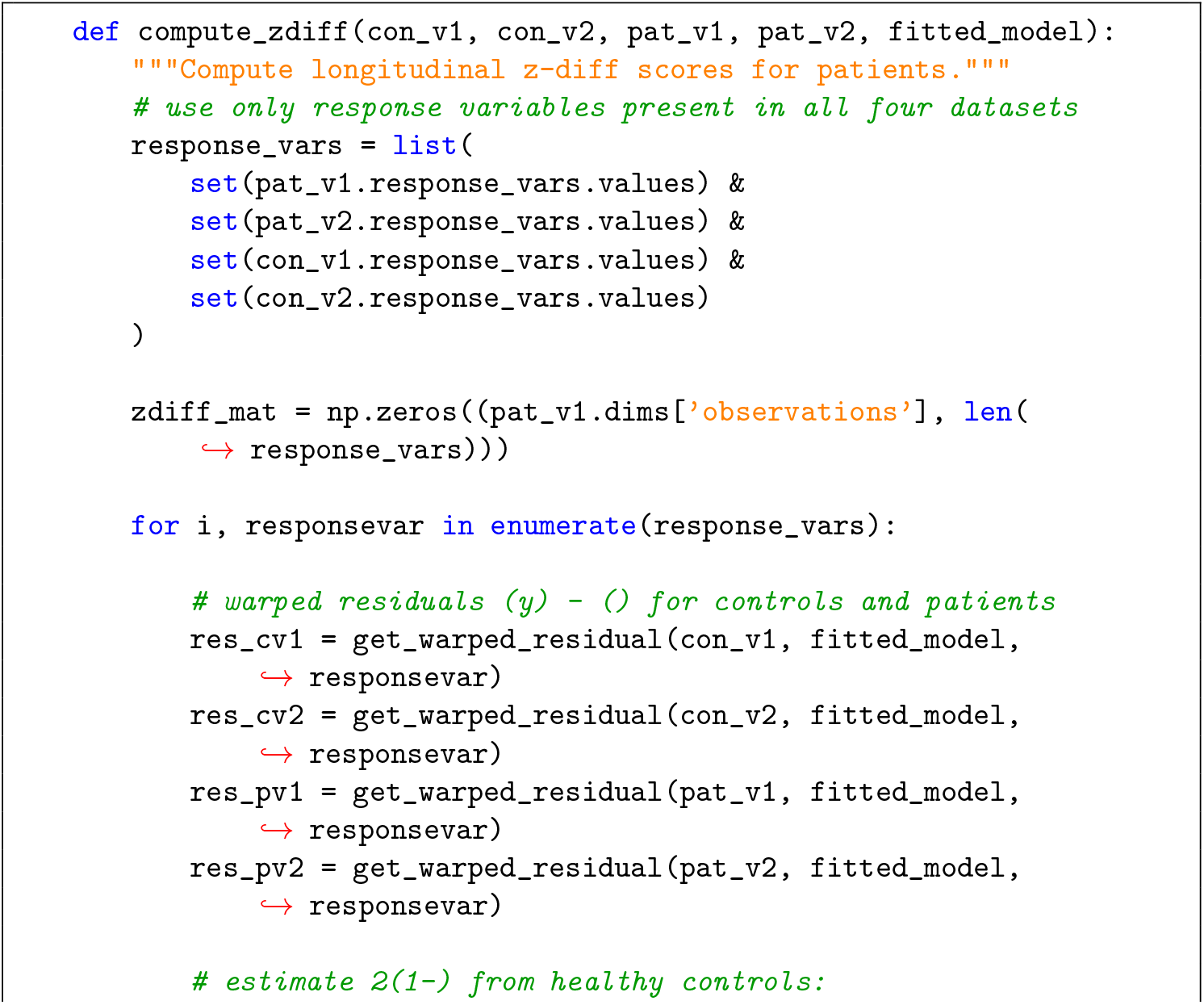
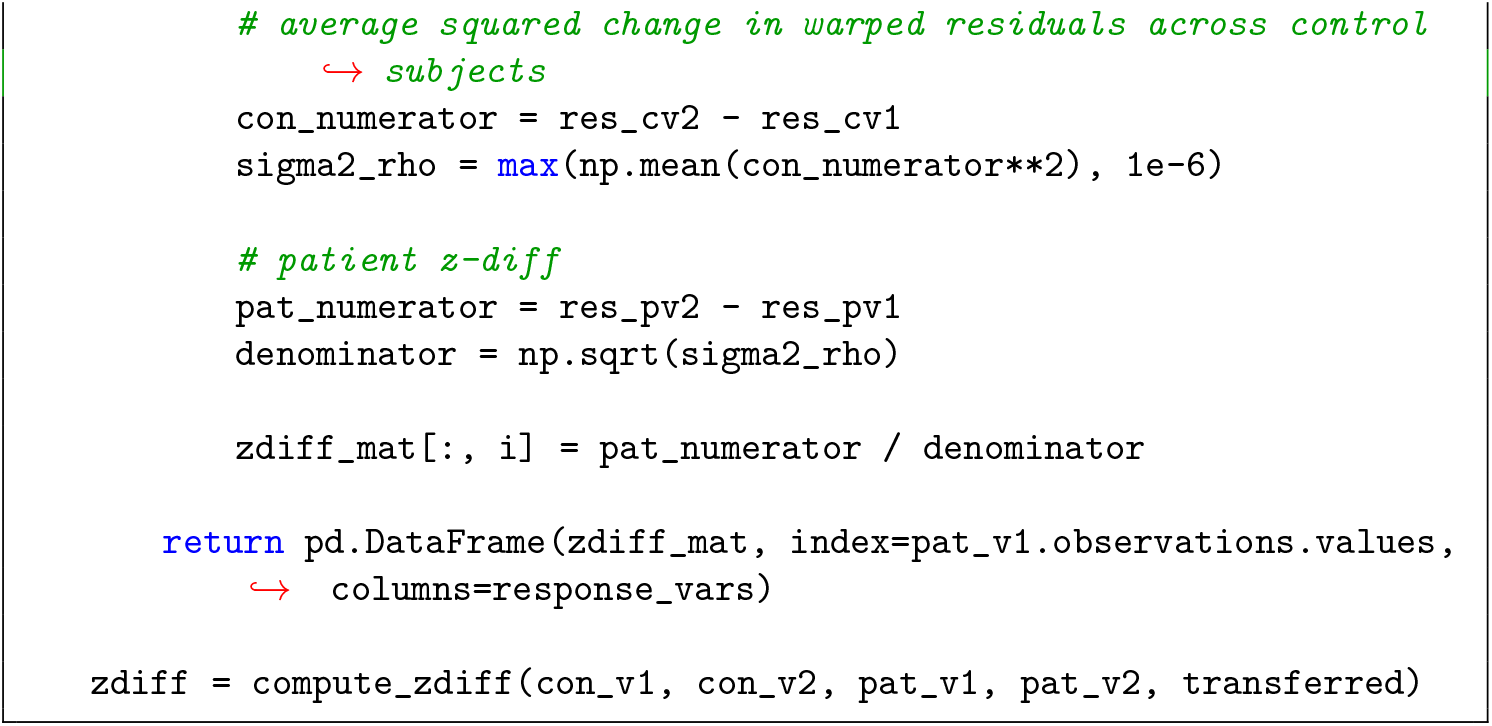
8. Evaluation: To identify brain regions with significant longitudinal change in patients, we apply a one-sample Wilcoxon signed-rank test to the z-diff scores for each response variable. This non-parametric test evaluates whether the median z-diff score across patients is significantly different from zero — i.e. whether patients show systematic deviation from the healthy ageing trajectory. To control for multiple comparisons across all response variables, we apply Benjamini-Hochberg (BH) false discovery rate (FDR) correction. 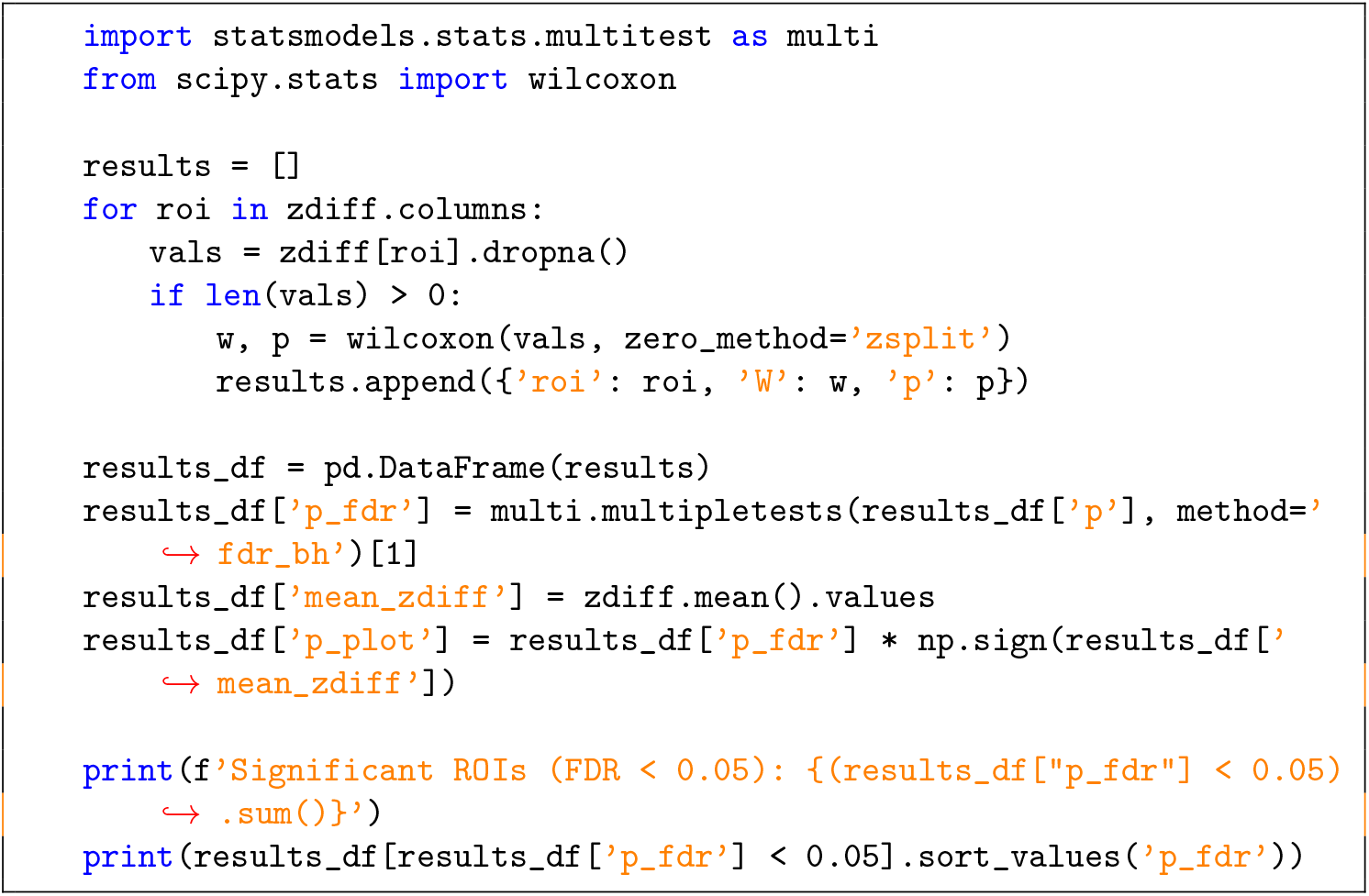

### Optional Workflow: Model Comparison TIMING ~5–7 min

Rank two (or more) fitted models, here the above-mentioned default HBR (10A) and custom HBR (10B), on their ability to predict unseen data using the ELPD criterion.

**PAUSE POINT:** This workflow provides a ranking for HBR models for the dataset at hand. For BLR, model comparison can be performed using the exact marginal likelihood by extracting the `nlZ` value from the fit `regression_model.json` file. However, it is important to recognise that whilst the marginal likelihood is obtained by integrating out the parameters (i.e. regression weights) does not automatically account for differences in the number of hyper-parameters (e.g. warp parameters), but these can be accounted for using the Bayesian information criterion. This can be computed using for `evaluate_bic` method in the NormData object. See [34] for details.

1. First, fit a default model to compare the main workflow model to 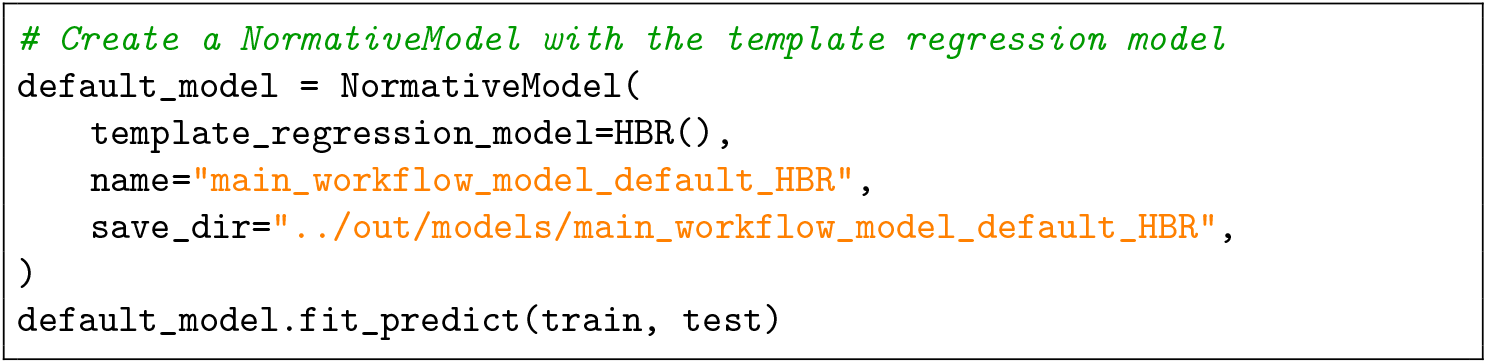
2. Run the model comparison 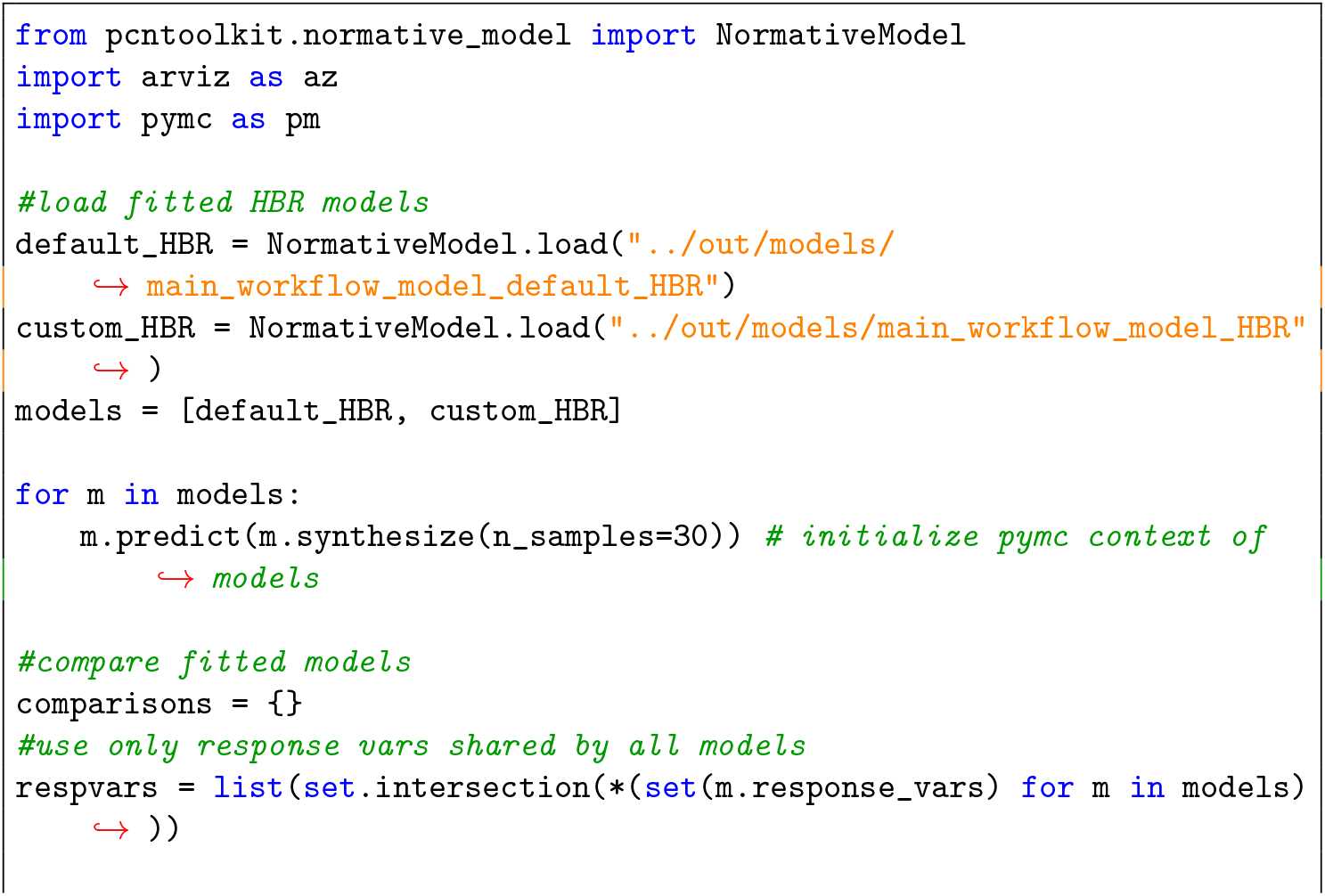
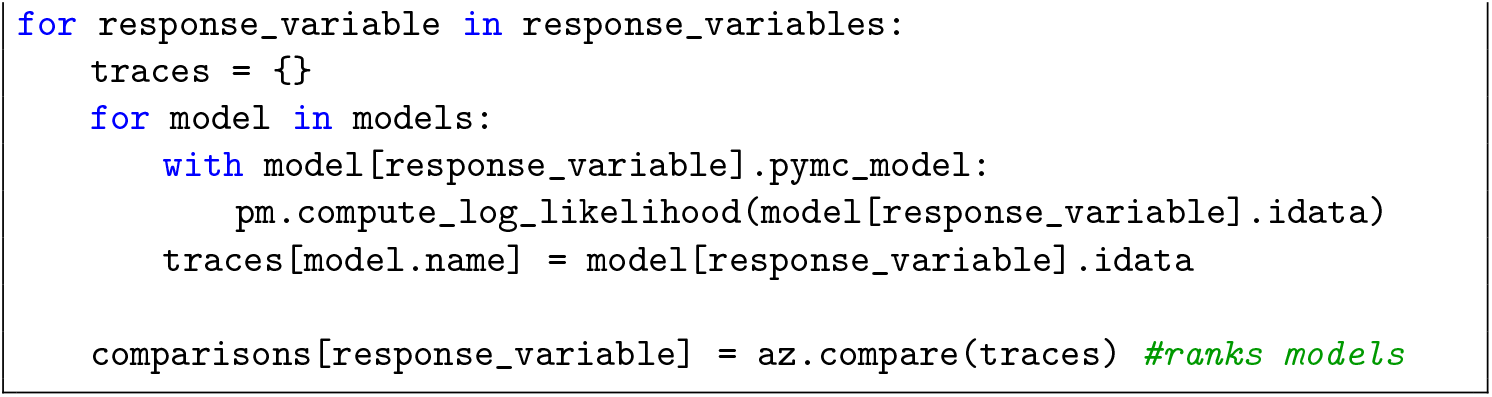

### Optional Workflow: Harmonization TIMING ~1–2 min

Remove group effects from data using the mapping between batch effect values and normative deviations values learned during model fit. This can be useful for example, to plot all data points harmonised on the same underlying centiles.

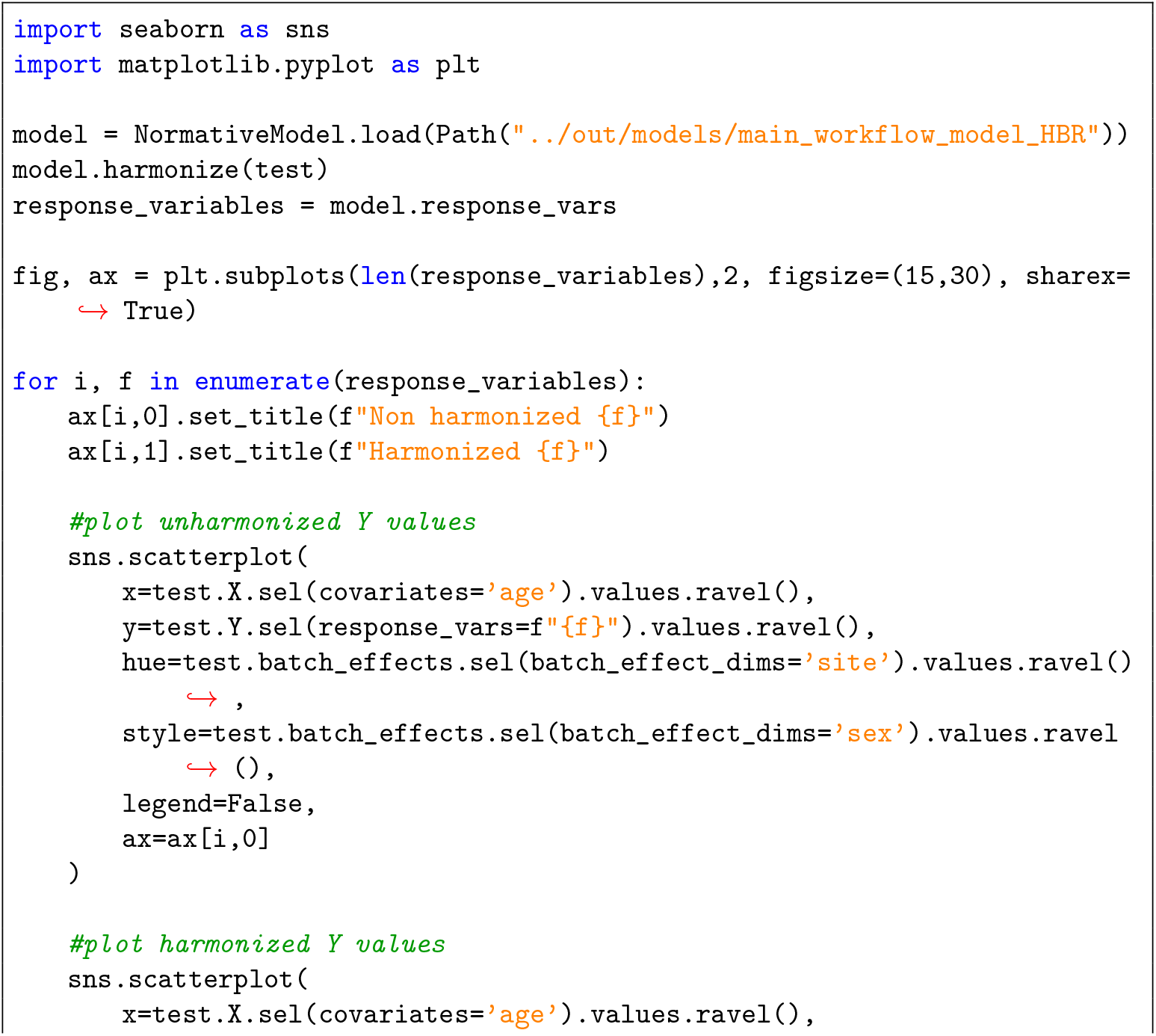
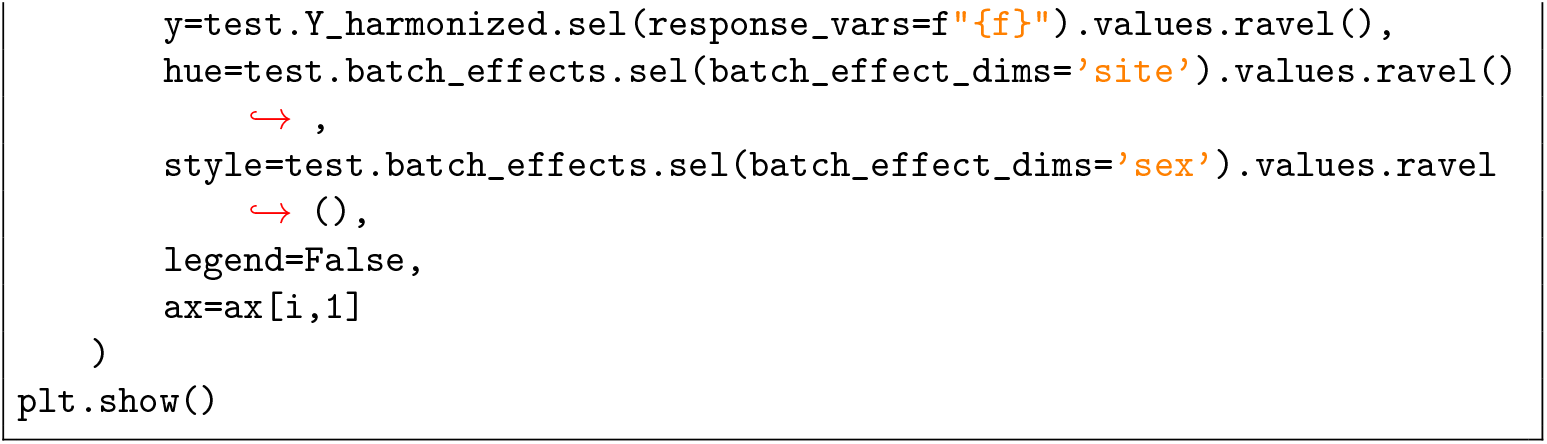

## 4 Anticipated Results

This protocol describes the process by which normative models can be fit. Starting from raw, multi-site neuroimaging data, we have described a reproducible end-to-end workflow covering data selection, preprocessing, model fitting, evaluation, and visualization including the assessment of model convergence and the fit of the centiles to the data. We have illustrated this process by providing a basic example of the main workflow using publicly available data. We elaborate on this in the next sections.

### An overview of model outputs

First we describe the low-level outputs from running the protocol. Running the workflow writes all intermediate and final outputs in the specified output path under `models/main_workflow_model_HBR/`. This directory contains the fitted model specifications, posterior samples, evaluation metrics, and visual diagnostics used in the subsequent sections.

- `model/` — per-feature subfolders containing a top-level `normative_model.json`, which is a serialized sumamry of the default configuration for the multi-feature model configuration (e.g. algorithm used, likelihood function, batch effect configuration etc). In addition, the software creates a separate folder for each feature, containing:
  – `regression_model.json`, serialized feature-level model specification. This contains regression model-specific information, such as the basis function configuration, the estimated means and variances and scaling parameters. As reflected in the protocol, this model inherits many of the attributes from the default configuration and provides a succint summary of all the information necessary to apply the normative model to new data (e.g. during a model transfer)
  – `idata.nc`: A netcdf file containing the posterior traces and diagnostics for HBR. This file is absent for BLR models.
- `plots/` — centile and quantile–quantile (QQ) diagnostics for both train and test sets, one set per feature.
- `results/` — CSV outputs with all numerical results:
  – `statistics {train,test} .csv`: fit metrics for each feature, including the root mean squared error (RMSE), explained variance (EXPV) and Shapiro-Wilks’ statistic. See ‘Interpretation of fit metrics’ for further details)
  – `centiles_{train,test`}`.csv`,
  – `Z_{train,test`}`.csv`,
  – `logp_{train,test`}`.csv`

A schematic of the default folder hierarchy:

~~~
models/
|-- main_workflow_model_HBR/
| |-- model/
| | |-- <feature_1>/
| | | |-- idata.nc
| | | ‘-- regression_model.json
| | |-- <feature_2>/
| | | |-- idata.nc
| | | ‘-- regression_model.json
| | ‘-- normative_model.json
| |
| |-- plots/
| | |-- centiles_<feature>_{train,test}_*.png
| | ‘-- qq_<feature>_{train,test}.png
| |
| ‘-- results/
| |-- centiles_{train,test}.csv
| |-- Z_{train,test}.csv
| |-- logp_{train,test}.csv
| ‘-- statistics_{train,test}.csv
~~~

Each folder is automatically generated by the PCNtoolkit during model fitting and prediction. Together, these outputs contain all information needed to reproduce the diagnostics, metrics, and visualizations, which are described in Model evaluation steps 12-14 of the Procedure. A fully commented walkthrough of the saving conventions, file contents, and plot interpretation is provided in the accompanying Jupyter notebook.

### Convergence diagnostics for example image derived phenotype

We show the results from one representative feature from the Freesurfer Destrieux atlas (*lh_G_and_S_frontomargin*) below.

Trace plots for the slope parameters of *µ* (`slope_mu`) are shown in Fig. 2. All four chains overlap tightly and explore the posterior space evenly, with no visual signs of non-stationarity, confirming that the hierarchical regression components have converged well.

### Interpretation of fit metrics

For this example, we evalute model performance using both mean-based and probabilistic metrics exported by the PCNtoolkit. Mean-based metrics including the root mean squared error (RMSE), standardized mean squared error (SMSE), and explained variance (EXPV) quantify how well the predicted mean *ŷ* approximates the observed data. As discussed in Section 1.5, probabilistic metrics such as the the mean standardized log loss (MSLL) and the mean absolute centile error (MACE) [44] assess the full predictive distribution; therefore, they are more suitable for evaluating the quality of estimated centiles in the NM context: MSLL compares predictive log-likelihoods to a variance-standardized baseline, while the MACE measures the average deviation of predicted centiles from their expected uniform targets.

Figure 3 shows the two plots produced by the code snippet in step 14 of the protocol. This shows a good fit in that the centiles align well with the data, evident both in the centile plots in panel (a) and in the QQ-plots in panel (b). For illustrative purposes, we also show an example of a poorly fitting model in Figure 4. These are derived from following the protocol outlined here for the white matter hypointensities phenotype derived from the Freesurfer subcortical parcellation for the FCON1000 dataset. This phenotype is very difficult to model because it is strictly non-negative, heteroskedastic and skewed. Accordingly, the simple Gaussian model shown in Figure 4(a-b) shows a poor fit. Whilst a Gaussian model can capture the heteroskedasticity and the non-linearity, it cannot capture the skew. This is reflected in the implausibly negative centiles (panel (a)) and the poor distributional fit of the evident in the QQ-plot (panel (b)). In contrast, a SHASH model provides a considerably better fit (Figure 4c-d) in that the centiles remain positive and the distributional fit is better in the qq-plot. Note that the fit is still not perfect, however, and a degree of misfit is evident in the extreme positive tail, probably driven by a few high leverage outliers also evident in the centile plot. This could be further improved by careful quality control, or by focussing analysis on the extreme tail of the distribution [45]. Whilst we do not cover how to fit SHASH models here, an additional workflow, providing a detailed didactic walkthrough for fitting SHASH models is provided in the associated [code repository](https://github.com/predictive-clinical-neuroscience/pu25_code/blob/main/notebooks/additional_worflows/HBR_SHASH_tutorial.ipynb).

**Fig. 4:**
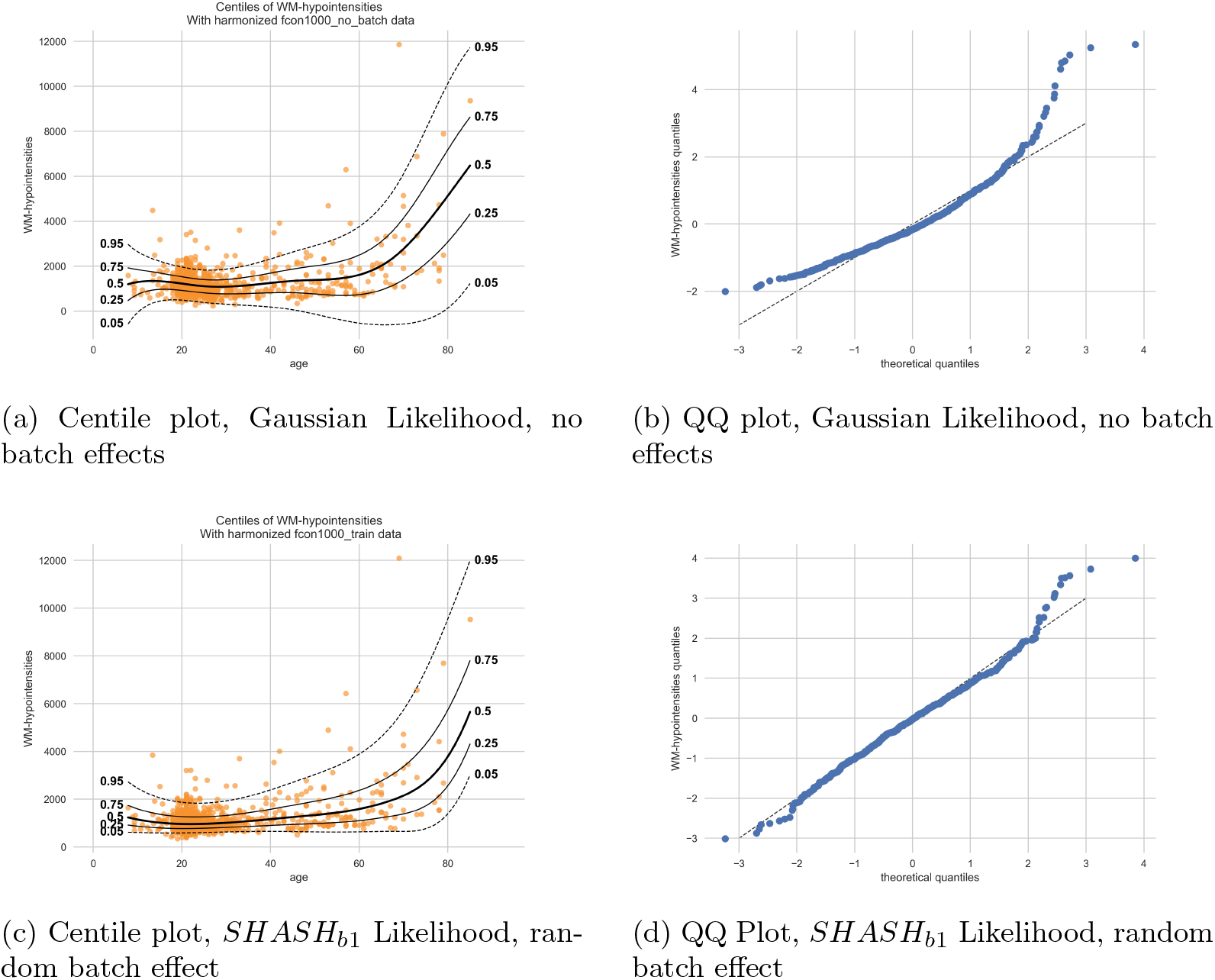
Visualization of good and bad model fit based on modelling choices in a non-gaussian, heteroscedastic phenotype. **(a)** The symmetry assumption of the Gaussian likelihood causes centiles to extend below zero and fails to capture the heteroskedasticity inherent in this phenotype. **(b)** This is reflected in the QQ-plot, where predicted and observed centiles diverge most strongly in age regions exhibiting non-Gaussian variance. **(c)** The lower centiles stay non-negative throughout, with asymmetric bands above and below the median. **(d)** The points lie much closer to the diagonal across the full range of centiles, with only minor deviations in the extreme tails.

In summary, for this example, these quantitative and visual diagnostics confirm robust model convergence, well-calibrated uncertainty estimates, and accurate functional dependence across covariates. The resulting normative model captures both central trends and population variability, allowing individual-level deviations to be expressed in a statistically calibrated manner.

### Optional federated learning

The outcome of the federated learning workflow is a set of statistics (Z statistics, fit metrics) that can be evaluated using the same tools as for the main workflow. Note that since the example here is based on BLR (not HBR), the assessment of model convergence is not necessary.

### Optional longitudinal modelling

The example workflow for the longitudinal normative modelling approach provides a set of ‘z-diff’ scores that can be used for downstream analysis. Since these are the same data that are reported in [11], we refer the reader to this publication for further information.

### Optional model comparison

Table 3 shows the expected outcome from the optional HBR model comparison (procedure section 17) that ranks fitted HBR models on their ability to generalize to unseen data (out-of-sample predictive fit), which is captured by the expected log pointwise predictive density (ELPD; the higher, the better)[19]. Models are ranked from best to worst, thus the custom model shown at rank 0 performs best in this example.

**Table 3:**
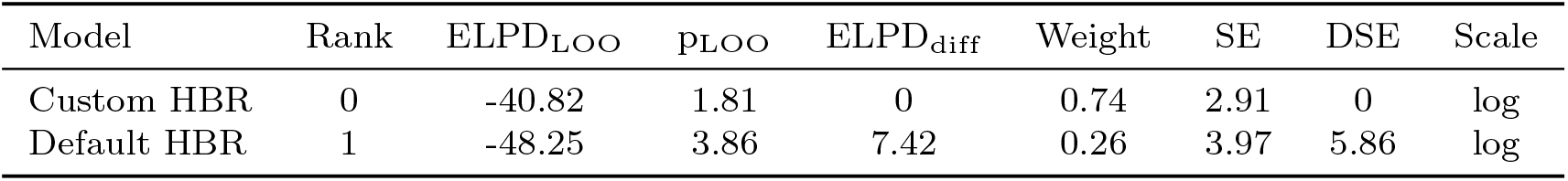
Best-to-worse ranking of fitted HBR models based on their out-of-sample predictive accuracy for an example feature (*lh G and S transv frontopol*). Default outputs include the expected log pointwise predictive density (ELPD) estimated with leave-one-out (LOO), p as the effective number of parameters (p_LOO_), the difference in ELPD between the top-ranking model and the others (ELPD_diff_), the relative weight for each model (probability of a model among other compared models given the data), the standard error of the ELPD (SE) and the standard error of the ELPD_diff_ (DSE). Log scale is used.

| Model | Rank | ELPD<sub>LOO</sub> | $p_{\text{LOO}}$ | ELPD<sub>diff</sub> | Weight | SE | DSE | Scale |
| --- | --- | --- | --- | --- | --- | --- | --- | --- |
| Custom HBR | 0 | -40.82 | 1.81 | 0 | 0.74 | 2.91 | 0 | log |
| Default HBR | 1 | -48.25 | 3.86 | 7.42 | 0.26 | 3.97 | 5.86 | log |

### Optional data harmonization

Figure 5 shows how feature values can be corrected for batch effects using a fitted normative model, following the optional workflow in procedure section 18. This is used by the centile plotting functions of the toolkit, but can also be used as an optional utility for downstream analyses, accounting for group effects in a test dataset without data leakage.

**Fig. 5:**
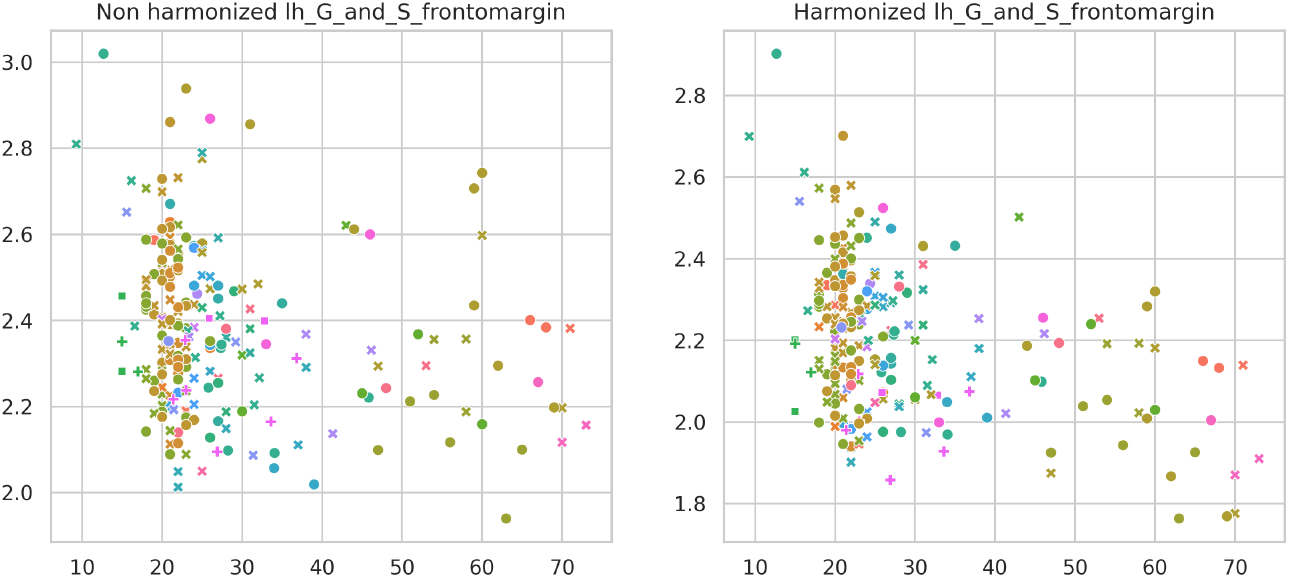
Visualization of example feature values (*lh G and S frontomargin*) before and after harmonization over batch effects. Values are harmonized over the first-ocurring unique combination of batch effects in the dataset.

## 5 Timing

The normative modeling portion of this protocol (including evaluation and visualization) can be completed in ~ 20 minutes. Most of the time is spent during the model configuration and fitting (steps 10-11, 5-15 minutes) and in the model evalation stages (steps 12-14, around 5-10 minutes). Steps 1-9 can be completed quickly (a few minutes). However, please note that the timings given are the time taken to actually run the code and evaluate the outputs. For a real analysis using different data, additional time will be required to evaluate different parameter settings, model configuration options etc. Please also note that these timing estimates are based on the use of the compute platform described above to run the code and the actual time will be dependent on the particular compute platform used.

## 6 Troubleshooting

A set of common problems the user may encounter during this protocol are listed in Table 4. Further, we have provided examples of a bad and good model fit in Figure 4.

**Table 4:** Common problems and solutions.

| step | problem | possible reason | solution |
| --- | --- | --- | --- |
| 2 | PyTensor warning: g++ not detected | C/C++ compiler is missing. HBR models will run very slowly without it (BLR is unaffected) | Install the C/C++ compiler via your terminal:<br>**Linux:** `sudo apt-get install g++`<br>**macOS:** `conda install -c conda-forge clangxx clang` |
| 3-8 | Unique values of a dimension in `NormData` do not match expected numbers | Formatting mismatch in data aggregation | Carefully check data types (int vs float vs str) before building `NormData`. |
| 11 | `KeyError` reporting that batch effect labels during model fitting | `make_compatible()` has not been run | Ensure `make_compatible()` is called *before* merging (step 7). |
| 11 | Compilation error during `fit_predict()` | C/C++ compiler is not installed or configured properly | Follow the installation instructions shown above for step 2 |
| 11 | File locking errors related to `hd5py` when running `fit_predict()` | File locking is enabled | Set the following environment variable:<br>`HDF5_USE_FILE_LOCKING=FALSE` |
| 14 | `KeyError` data and centiles do not align, for example when plotting the data | The data and centiles are plotted for different batch effect combinations | Ensure a `reference_batch_effect` is specified and all data are harmonized to the same space. |
| 14 | Centiles assume implausible values (e.g. negative values for volumes) | A continuous model may have been used for strictly positive data | Consider specifying `y_transform="log"` as a keyword argument in `NormativeModel()` to enforce positivity. |
| 14 | Centiles show poor calibration | Chosen likelihood function may not be appropriate for the data | Consider using a different likelihood function. |

Further resources to assist troubleshooting are provided in the additional documentation available online at [ReadTheDocs](https://pcntoolkit.readthedocs.io/en/latest), including additional tutorials for other algorithm implementations, a glossary to clarify the jargon associated with the software, a reference guide with links to normative modeling publications and a frequently-asked-questions page where many common errors (and their solutions) are discussed in detail. Additionally, questions can be raised and answered by submitting issues via GitHub.

## 7 Key references

- De Boer et al. [9]
- Bayer et al. [12]
- Fraza et al. [46]
- Berthet et al. [47]
- Cirstian et al. [18]

## Acknowledgments

We wish to acknowledge the contributions of the many users of our toolbox who have provided feedback or made suggestions that have enabled us to improve our software.

## Author Contributions

AAAdB, JMMB, CF, AC, BRB: writing software, running experiments, drafting and revising the manuscript; AC: running experiments, revising the manuscript; KT, ES: writing software, drafting and revising the manuscript AB, RC, MZ, SR, AAK: running experiments, drafting and revising the manuscript; TW, CFB, SMK: conceptualisation, supervision, drafting and revising the manuscript; AFM: funding acqusition, conceptualisation, writing software, supervision, drafting and revising the manuscript.

## Data Availability

All data used in this manuscript are publicly available. This includes the FCON1000 dataset, which is freely available at the [FCON1000 website](https://fcon_1000.projects.nitrc.org/indi/fcp_abcd/abcd.html) and four clinical datasets from the OpenNeuro portal, namely:

1. ds003568, see ref: [39].
2. ds003653, see ref: [40].
3. ds003469, see ref: [41]
4. ds000222, see ref: [42]

For the longitudinal workflow, we use an additional clinical dataset, see ref: [11]. These (pseudonymised) data are provided in the associated code repository.

## Code Availability

The PCNtoolkit software package is available online at : https://github.com/amarquand/PCNtoolkit and all code necessary to run this protocol is available at: https://github.com/predictive-clinical-neuroscience/pu25_code.

## Footnotes

1 Since there are extensive changes in underlying data storage formats and object oriented structure, we recommend that users of the software re-fit all models fit using versions of PCNtoolkit prior to `1.x.x`. Since this version of the toolkit was designed with longevity in mind, we do not envisage future deprecations for the model structure.

2 Note that these diagnostics apply specifically to HBR models, which rely on Markov Chain Monte Carlo sampling. For BLR models, estimated via optimization, different criteria apply, typically based on gradient norms and optimizer tolerance. However the optimisation framework used in PCNtoolkit is fairly robust to these differences, so it is not normally necessary to check these for BLR models.

